# Functional Inference of Gene Regulation using Single-Cell Multi-Omics

**DOI:** 10.1101/2021.07.28.453784

**Authors:** Vinay K. Kartha, Fabiana M. Duarte, Yan Hu, Sai Ma, Jennifer G. Chew, Caleb A. Lareau, Andrew Earl, Zach D. Burkett, Andrew S. Kohlway, Ronald Lebofsky, Jason D. Buenrostro

**Affiliations:** Department of Stem Cell and Regenerative Biology, Harvard University, Cambridge, MA 02138, USA; Gene Regulation Observatory, Broad Institute of MIT and Harvard, Cambridge, MA 02142, USA; Bio-Rad, Digital Biology Group, Pleasanton, CA 94588, USA; Department of Pathology, Stanford University School of Medicine, Stanford, CA 94305, USA

## Abstract

Cells require coordinated control over gene expression when responding to environmental stimuli. Here, we apply scATAC-seq and scRNA-seq in resting and stimulated human blood cells. Collectively, we generate ∼91,000 single-cell profiles, allowing us to probe the *cis*-regulatory landscape of immunological response across cell types, stimuli and time. Advancing tools to integrate multi-omic data, we develop FigR - a framework to computationally pair scATAC-seq with scRNA-seq cells, connect distal *cis*-regulatory elements to genes, and infer gene regulatory networks (GRNs) to identify candidate TF regulators. Utilizing these paired multi-omic data, we define Domains of Regulatory Chromatin (DORCs) of immune stimulation and find that cells alter chromatin accessibility prior to production of gene expression at time scales of minutes. Further, the construction of the stimulation GRN elucidates TF activity at disease-associated DORCs. Overall, FigR enables the elucidation of regulatory interactions across single-cell data, providing new opportunities to understand the function of cells within tissues.

## Introduction

Eukaryotic cells have evolved exquisite control to continuously sense and respond to external environmental cues^1–4^. This, in part, involves coordinated changes in signaling dynamics, transcription factor (TF) binding, and eventually the expression of downstream target genes^3–5^. Immune cells, in particular, harbor tremendous plasticity in their ability to respond to stimuli, developing both diverse and specific functions in response to different pathogenic agents^6^. This highly context-specific, and often heterogeneous, activation of genes promoting the appropriate anti-viral or inflammatory response comprises one of the hallmarks of immunity. Our understanding of immunity has evolved over time, for example it has been shown that chromatin may prime cells for immunological response^7, 8^, leading to exhausted states^9^ or further orchestrating the activation of surrounding cells through the production of key signaling molecules^10^.

Single-cell genomics methods have greatly advanced our understanding of cellular diversity of immune cells^11–13^. Single-cell RNA-sequencing (scRNA-seq) characterizing time- and stimulus-dependent transcriptional signatures in mouse^10^ and human^14^ immune cells, for example, have identified distinct transcriptional programs that are activated or repressed over time and highlighted cell-cell variability in response to immunological stimulants^15^. Concomitantly, several prior studies have applied chromatin accessibility and gene expression assays to define *cis*-regulatory atlases across resting^12, 16, 17^ and stimulated^14, 18^ immune cell types. Most recently, the COVID pandemic has further prompted the use of single-cell ATAC-seq and RNA-seq tools to characterize the immunological response to infection^19, 20^. These diverse efforts have all sought to elucidate the epigenetic control of immune cell function, namely the cellular circuitry that defines the gene regulatory network (GRN) within the cell.

While these efforts have resulted in tremendous insights into the transcriptional control of immune cells, collectively these studies are limited by the existing repertoire of computational tools modelling gene regulatory dynamics among single cells. Recent advances in constructing GRNs from single-cell data^21, 22^ have facilitated new opportunities to uncover mechanisms of cell function and adaptation following stimulus. However, approaches that solely utilize co-expression^19, 20^ are limited in their ability to determine the presence of (i) master TF regulators and (ii) key *cis*-regulatory elements that activate the expression of genes. To this end, extensive prior work has demonstrated that epigenomics data can vastly improve the determination of functional GRNs^23–25^. Thus, we posit that computational methods for building GRNs that leverage high-throughput single-cell multi-omic data (ATAC and RNA) would improve our understanding of the epigenetic mechanisms underlying the function and adaptation upon environmental exposures of eukaryotic cells.

Here, we create an exemplar dataset for the construction of immune cell GRNs. To do this, we combine the use of multiple stimulus agents together with chromatin accessibility and gene expression single cell analysis to characterize and assess the dynamics of the *cis*-regulatory landscape linked with immune cell stimulation in human peripheral blood mononuclear cells (PBMCs). We then establish Functional inference of Gene Regulation (FigR), an generalizable approach for independently or concomitantly-profiled single-cell ATAC-seq (scATAC-seq) and scRNA-seq, that i) computationally-pairs scATAC-seq and scRNA-seq datasets (if needed), ii) infers *cis*-regulatory interactions, and iii) defines a TF-gene GRN. Utilizing these integrated data, we establish that changes in chromatin accessibility foreshadows changes in gene expression upon immune stimulation of monocytes. Last, we highlight how this approach can be used to identify key TFs and their relationship to target genes, including stimulus response and disease-associated Domains of Regulatory Chromatin (DORCs). Collectively, our work highlights the use of blood stimulation combined with high-throughput single-cell multi-omics, and advancements in developing GRNs using FigR, as a model to deduce key transcriptional regulatory modules that are required for immune cell activation.

## Results

### Combined high-throughput single cell epigenomic and transcriptional profiling of resting and stimulated PBMCs

To characterize the chromatin accessibility and transcriptional landscape associated with host response to stimuli in human blood, we performed droplet-based single-cell ATAC-seq (scATAC-seq) and single-cell RNA-seq (scRNA-seq) on both resting and stimulated human PBMCs for different time points of stimulus exposure (**Fig 1A**; see Methods). Specifically, cells derived from healthy donors (*n* = 3 or 4; **Table S1**) were exposed for either 1 or 6 hours (h) to stimulants known to elicit anti-viral-like or core inflammatory responses, including lipopolysaccharide (LPS) - a component of bacterial cell membranes, phorbol myristate acetate (PMA) plus ionomycin - a potent ester that activates NF-kB signaling^26^, or interferon gamma (IFN-γ) - an endogenously-produced immunoregulatory cytokine, alongside a DMSO control per time point, prior to single cell profiling (see Methods). These stimulants were chosen as they have been shown to induce distinct time and cell type-specific changes with unique transcriptional dynamics as part of the host immune response^10, 14, 26–28^. Additionally, for the 6-hour time point using each stimulant, we separately treated cells with a Brefeldin A, a protein secretion inhibitor (Golgi Inhibitor, GI), hence attenuating paracrine signaling events in immune cells allowing us to distinguish between primary versus secondary stimulation response phenotypes.

**Figure 1.**
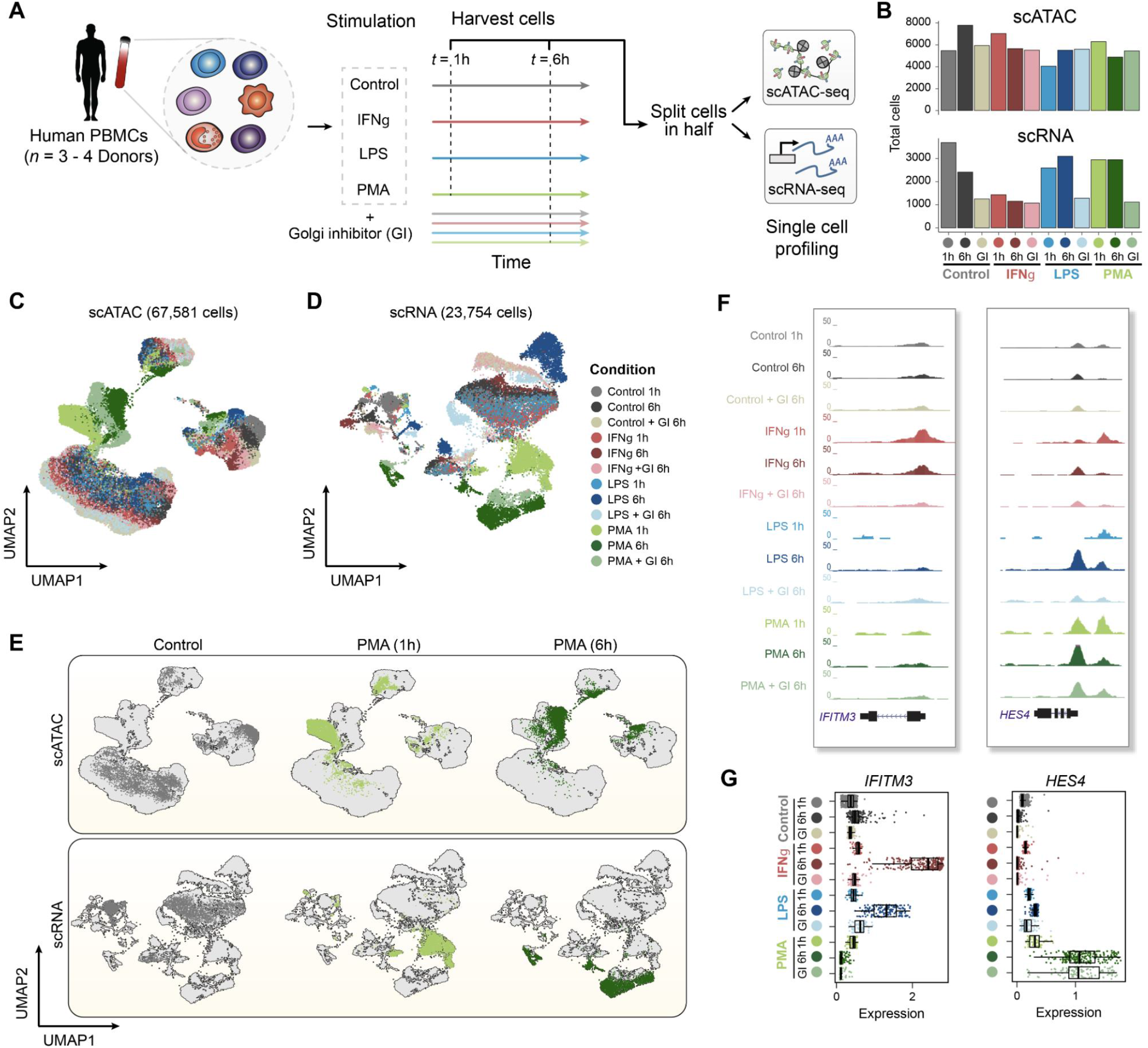
High-throughput single-cell epigenomic and transcriptional profiling of resting and stimulated human blood cells. **A.** Schematic highlighting design of stimulation experiment. Human peripheral blood mononuclear cells (PBMCs) were stimulated with either DMSO control, lipopolysaccharide (LPS), interferon gamma (IFN-γ) or phorbol myristate acetate plus ionomycin (PMA) for 1 hour or 6 hours, with or without a Golgi inhibitor (GI) for the 6-hour treatment condition. Cells were then split and profiled using scATAC-seq and scRNA-seq for each condition and time point considered. **B.** Total number of cells profiled per condition passing quality control filtering for scATAC and scRNA-seq. **C.** UMAP projection of scATAC-seq cells based on LSI dimensionality reduction, with cells colored by treatment condition. **D.** UMAP projection of scRNA-seq cells based on PCA dimensionality reduction, with cells colored by treatment condition. **E**. UMAPs of scATAC-seq cells (top) and scRNA-seq cells (bottom) highlighting individual conditions in control (6h) and PMA (1h and 6h) conditions. **F.** Aggregate accessibility profiles for scATAC-seq monocyte cells around genes *IFITM3* and *HES4*. **G.** Distribution of single cell expression levels based on the imputed scRNA-seq counts for stimulation-specific gene markers shown in F per condition for scRNA-seq monocyte cells.

Collectively, we generated over 15 billion reads resulting in a high-coverage single cell regulatory atlas comprising of 67,581 scATAC-seq and 23,754 scRNA-seq cells spanning all conditions (**Fig 1B**), with an average of 8,865.2 (±s.d.= 4,837) aligned unique nuclear fragments per cell and mean fraction of reads within peaks (FRiP) of 0.6 (±s.d. = 0.05) for scATAC-seq profiled cells (**Fig S1A-B**), and averaging 3,021 UMIs (±s.d. = 425.77) for scRNA-seq profiled cells (**Fig S1C-D**; See Methods). Clustering scATAC-seq and scRNA-seq cells (see Methods) yielded discrete cell clusters, largely representing monocytes, T (CD4/CD8) and B lymphocytes, and natural killer (NK) cells, with even distribution of cells from all donors involved per cluster and condition (**Fig1C-D** and **Fig S1E-G**). Importantly, each of these broader clusters included sub-clustering of cells by stimulus condition (**Fig 1E** and **S1H**).

To formally annotate cell types for scRNA-seq cells, we first aligned cells across batches (here, defined as each treatment condition) using a previously described computational approach^29^, enabling the co-clustering and annotation of scRNA-seq cells across conditions. Clustering of cells using this approach yielded distinct groupings (**Fig S2A-B**), which were enriched for cell type and stimulus-specific gene expression markers and were used to annotate cell types (**Fig S2C**). Furthermore, inspection of the myeloid cells for accessibility peaks around gene promoters (scATAC-seq), and gene expression levels (scRNA-seq) confirmed stimulus and time specific changes (**Fig 1F-G; Fig S2D**). Importantly, all major cell types were captured at relatively even proportions across the treatment conditions used (**Fig S2E**), enabling multi-omic integration of independently assayed chromatin accessibility and gene expression profiles downstream.

### A computational cell pairing approach for accurate integration of single-cell chromatin accessibility and gene expression profiles

We reasoned that data from paired contexts may enable the determination of gene regulatory networks (GRNs), facilitating the interpretation of the key regulatory processes underlying stimulus of immune cells. Current frameworks supporting the integration of scATAC and scRNA-seq data^29–31^ rely on identifying ‘anchor’ cells - cells that represent shared biological states in a common lower dimensional space, to then find representative cells from one dataset in the other. While useful for matching cells of corresponding cell types (i.e. annotation-level pairing), these methods often i) result in high one-to-many cell barcode matching rates, resulting in overall lower cell usage downstream, or ii) do not adequately address cell type imbalance between datasets.

To address this challenge, we developed a method (OptMatch) that identifies cell pairs between scATAC-seq and scRNA-seq data using a constrained optimal cell mapping approach (**Fig 2A**). For this approach, we first create a shared co-embedding of scATAC-seq and scRNA-seq cells using canonical correlation analysis (CCA), similar to what has been previously described^29^. Next, we address the issues of i) total cell number imbalance and ii) cell type imbalance between datasets by first sub-clustering the entire cell space and constructing a cell *k*-nearest neighbor (kNN) graph between ATAC and RNA cells in the co-embedded space, sampling cells from both assays within a given kNN subgraph (see Methods). Upon down-sampling to match cell numbers between assays (i.e. scATAC or scRNA) in a given subgraph, cells are then paired using a constrained global matching algorithm^32^, using the subgraph geodesic distance between ATAC-RNA cells as a cost function. Analogous to the traveling salesman problem, this ensures that resulting ATAC-RNA cell pairs are minimized for the total geodesic distance among all combinations of possible pairs. Importantly, only ATAC-RNA cells within a certain distance (geodesic kNNs) are considered for pairing as a prior, further speeding up computation time relative to if all possible pairs were being considered (see Methods).

**Figure 2.**
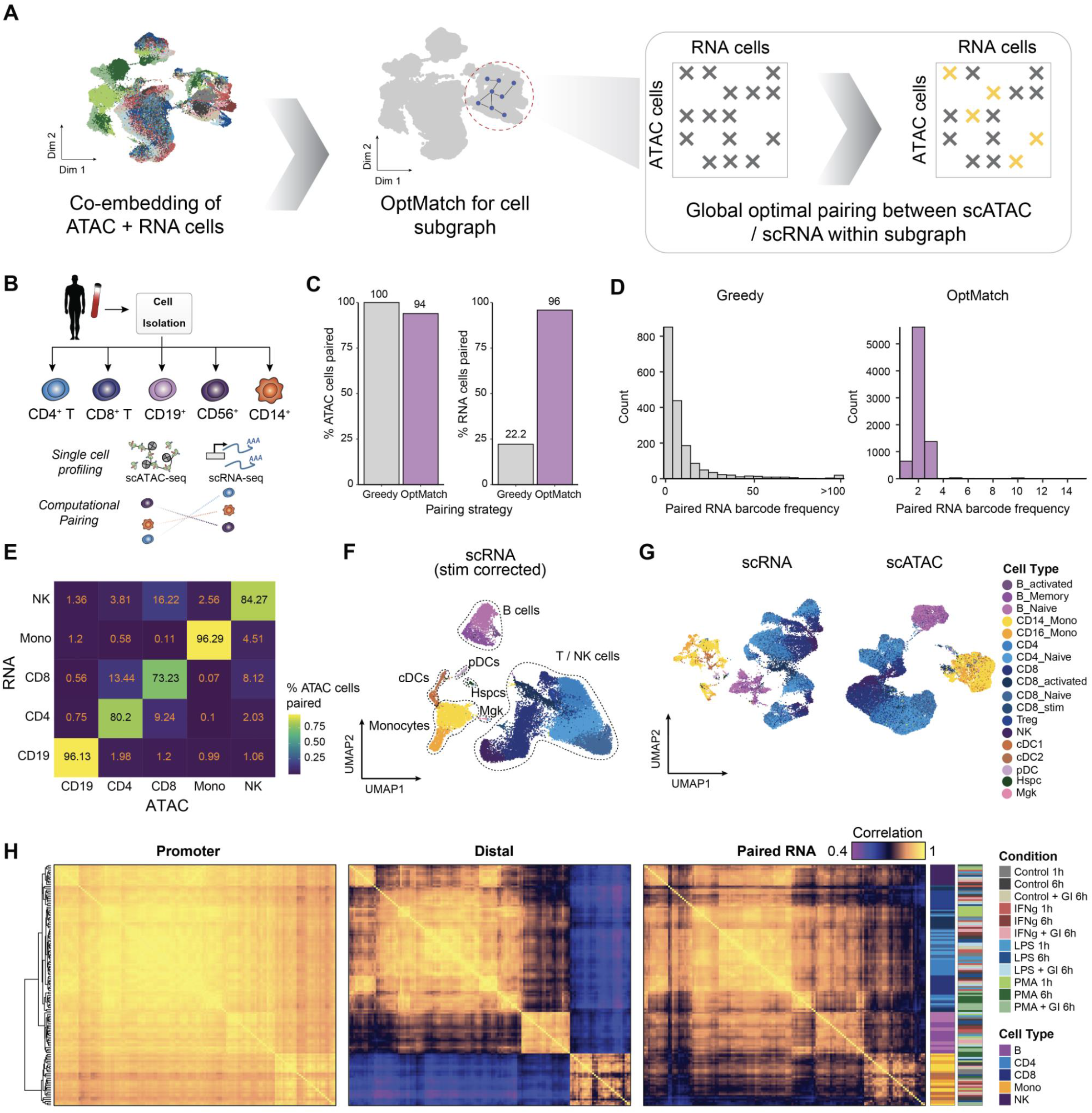
Sparse kNN-based ATAC-RNA cell pairing allows optimal pairing and integration of scATAC-seq and scRNA-seq data. **A.** Schematic highlighting strategy for computational pairing of scATAC-seq and scRNA-seq cells based on geodesic distance *k*-nearest neighbors (yellow x marks) within cluster subgraphs (gray x marks). **B.** Schematic depicting experimental bead enrichment of specific immune cell types from human PBMCs. **C.** Distribution of the number of instances of paired RNA cell barcode when using the greedy (left) versus OptMatch method for the PBMC isolate dataset pairing. **D.** Percentage of total scATAC and scRNA-seq cells paired using the two different pairing strategies. UMAP projection of bead-enriched PBMCs profiled using scATAC-seq (left) and scRNA-seq (right). scATAC-seq cells are clustered based on peak accessibility, while scRNA-seq cells are clustered based on variable gene expression. **E.** Accuracy heatmap of scATAC-scRNA-seq pairing between PBMC isolate cell types, colored by percentage of scATAC-seq cells correctly paired to the corresponding scRNA-seq cell type. **F.** UMAP of scRNA-seq stimulated cells shown in Fig 1D, with cells aligned across stimulus conditions to enable cell type annotation, colored by annotated cell type **G.** UMAP of un-aligned scRNA-seq cells (shown in Fig 1D) colored by annotate cell type, and scATAC-seq stimulated cells (shown in Fig 1C), colored by paired scRNA-seq cell annotations, enabling downstream data integration for stimulated scATAC and scRNA-seq profiled cells. **H.** Pairwise Pearson correlation of aggregate single cell chromatin accessibility profiles associated with gene promoters (left), distal from the promoter (center) and paired gene expression (right), aggregated by cell type and condition.

To create a reference data set to benchmark OptMatch, we isolated cell types within PBMCs and profiled (in separate assays) scRNA-seq and scATAC-seq^13^. The complete data reflected scATAC-seq (*n*=17,920 cells) and scRNA-seq (*n*=8,089 cells) data corresponding to five PBMC sub-populations (**Fig 2B** and **Fig S3A-F;** see Methods). Using this data, we determined ATAC-RNA cell pairs using either: i) the optimal matching described above (OptMatch) or ii) a “greedy” best match approach (choosing the closest RNA cell for every ATAC cell in CCA space). As expected, we found OptMatch results in a significantly larger number of cells being paired from both datasets across all cells (92.06% scATAC and 98.4% scRNA; **Fig 2C** and **Fig S3G**), a consequence of fewer ATAC-RNA cell multi-mapping instances (**Fig 2D** **and Fig S3H**) compared to the greedy approach (22.2% scRNA). Importantly, the OptMatch approach also accurately maps cells of the same reference cell type (**Fig 2E**).

Motivated by the high accuracy of the OptMatch pairing approach, we sought to apply it to pair our stimulus multi-omic datasets. Pairing of scATAC and scRNA-seq cells per condition using OptMatch (**Fig S3I-K**), we obtained paired multi-omic data with matching cell numbers across assays (*n*=62,219). Importantly, this cell pairing further enabled cell type annotation of scATAC-seq by simply using annotations defined from scRNA-seq gene expression markers (**Fig 2F-G** and **Fig S3L**). Aggregating single cells by cell type and condition, and filtering for sufficient counts, resulted in 139 pseudobulks (averaging a total of 1.94M RNA and 2.3M ATAC aggregate counts). Utilizing this high-depth resource, we find that chromatin accessibility at distal peaks are highly cell type specific, even more so than gene expression, whereas promoter accessibility is relatively invariant across cell types and stimulation conditions (**Fig 2H**), validating prior reports^17, 33^. Overall, the high quality of this data and exquisite cell type specificity of distal chromatin accessibility motivated further analysis into the gene regulatory network underlying stimulus response. To this end, we reasoned that this OptMatch approach for cell pairing, enabling approximately uniform pairing of scATAC to scRNA profiles, would establish an integrated data set and could be used for downstream analysis analogously to accessibility and RNA expression profiling concomitantly within the same cell.

### Identification of distal peak-gene interactions across stimulation using integrated single cell data

We next sought to associate changes in *cis*-regulatory peaks to the expression of genes as a means to prioritize features that are part of the immunological response GRN. To do so, we built upon a previously described framework to establish significant distal peak-to-gene expression interactions^34^. Specifically, we used computationally paired cells (*n*=62,219 cells per assay) to correlate accessibility from peaks found within a fixed window (100 kb) around each gene’s transcription start site (TSS) to the expression of that gene, with permutation-based testing to estimate the statistical significance for a given peak-gene pair (**Fig 3A**; See Methods). In this way, we identified a total of 34,370 unique chromatin accessibility peaks genome-wide showing a significant association to gene expression (permutation *P ≤* 0.05), spanning a total of 11,304 genes. Prioritizing genes based on their total number of significantly correlated peaks, we identified a subset of genes associated with a high-density of peak-gene interactions, which we recently described as domains of regulatory chromatin (DORCs)^34^ (**Fig 3B**; *n ≥* 7 significant peak-gene associations; *n*=1,128 genes; *n*=12,583 peaks).

**Figure 3.**
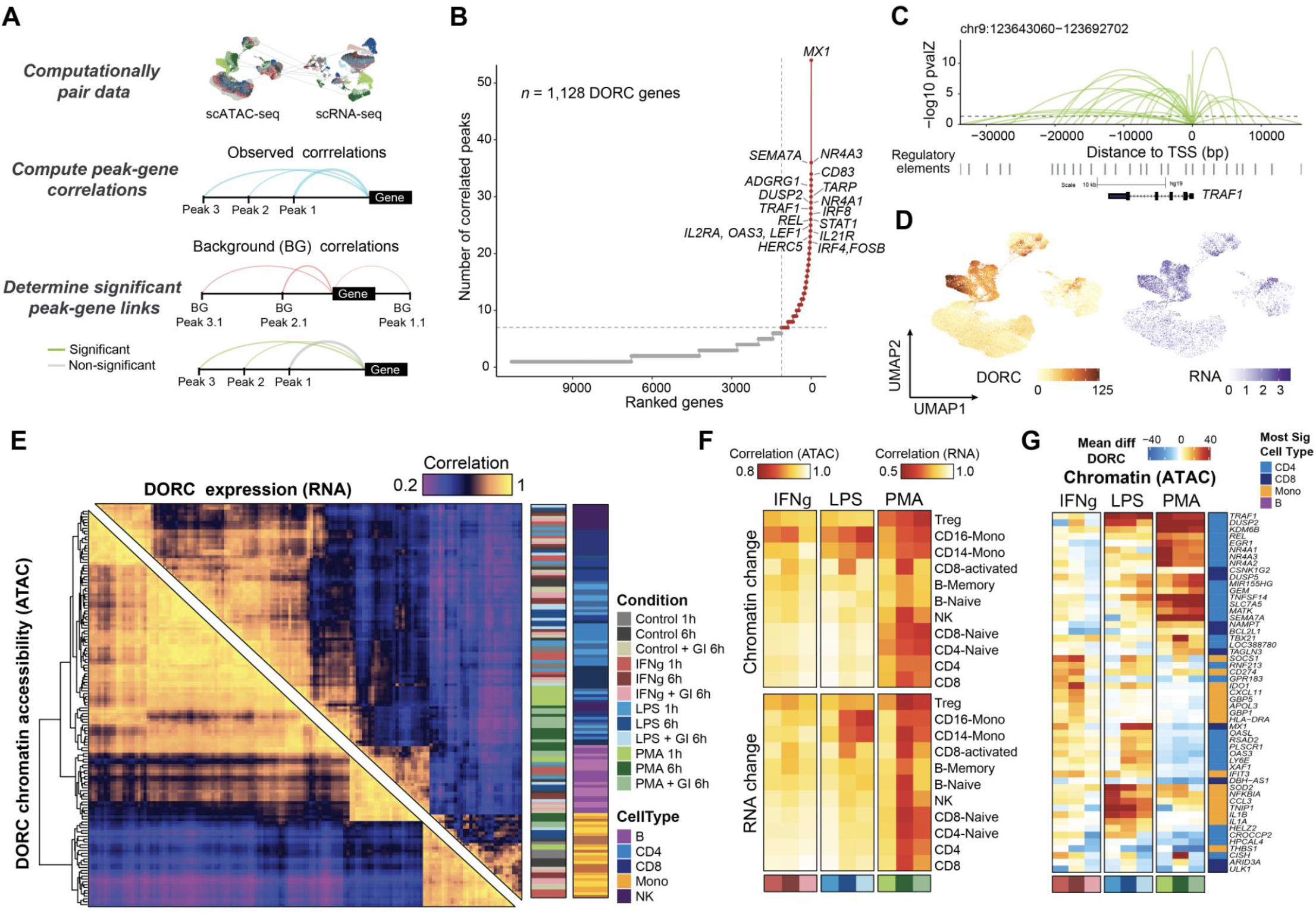
Integrative multi-omic analysis identifies key regulatory modules associated with stimulus response in single cells. **A.** Schematic of *cis*-regulatory analyses for identification of significant chromatin accessibility peak-gene associations using computationally-paired scATAC-seq and scRNA-seq stimulation datasets. **B.** Top hits based on the number of significant gene-peak correlations across all cell types and stimulus conditions. **C.** Loop plots highlighting significant peak-gene associations for DORC *TRAF1*, determined using the approach outlined in A **D.** UMAP of DORC accessibility scores (left) and paired RNA expression (right) for *TRAF1*. **E.** Pairwise Pearson correlation of aggregate DORC accessibility scores and RNA expression of cells per condition per cell type across all DORCs, clustered using hierarchical clustering by DORC score correlations. **F.** Global DORC accessibility (top) or gene expression (bottom) change displayed based on the Pearson correlation coefficient of the aggregate score across DORCs for each stimulation condition vs its corresponding control condition, shown per condition per cell type annotation. **G.** Heatmap showing the mean difference in single cell DORC accessibility, for the union of the top 10 differential DORCs across conditions and cell types (*n*=53 genes). Cell type color bar represents the cell group having the most significant change across all conditions, for that assay.

The list of DORC-associated genes included many known mediators of immunological response associated with innate and adaptive immune response pathways^10, 27, 35, 36^, as also confirmed by gene set enrichment analysis (GSEA) (**Fig S4A** and). Notably, among these genes, we see a large fraction of distal *cis*-regulatory associations (>5 kb away from the gene TSS; **Fig 3C** and **Fig S4B-C**). By scoring cells using the total associated peak accessibility signal per DORC (referred to as the DORC accessibility score), we determine correspondence between chromatin accessibility and gene expression across single cells (**Fig 3D** and **S4D-E**) or across pseudobulks for each DORC, stimulation condition and cell type (**Fig 3E**). Upon comparison with matched control conditions (DMSO controls), we observed the largest effect on DORC accessibility and expression from treatment with PMA, as seen across most cell types, and a more moderate effect with treatment of IFNγ or LPS, as seen predominantly in monocytes (**Fig 3F**). Notably, we found that stimulation induces a larger change in the transcriptome of the cells, in comparison to chromatin accessibility, however cell types concordantly altered both chromatin and expression to induce activation of immunity genes (**Fig 3E-F**). Interestingly, we also find that the addition of the Golgi inhibitor strongly attenuates immune response to PMA (CD8, NK and B) and LPS (CD8) - likely a consequence of inhibiting paracrine signaling - and in response to IFN (monocytes) - likely a consequence of inhibiting autocrine signaling.

Single cell differential testing among DORCs identified a number of essential regulators of immunological response (**Fig 3G** and **Fig S4F**). This includes shared LPS and IFN-induced genes (*MX1*, *IFIT3*, *OAS3*, *OASL*), and PMA-induced genes associated with cellular apoptosis and survival (*NR4A1/2/3, EGR1, REL, TRAF1*). Interestingly, we also observe primary ligand-encoding genes (*IL1A*, *IL1B, CCL3*) and immune inhibitors (*CD274* - also known as *PDL1*, *NFKBIA*, *TNIP1*) among these top differential DORCs. Notably, our *cis*-regulatory analysis recovers DORCs the majority of which (∼79%) include genes previously annotated to be linked to super-enhancer regions across diverse cellular contexts (See Methods) (**Fig S4G-H**), the remainder (*n*=238 genes) including several stimulation-response genes (*IFIT1*, *MX1, OAS1/3, IL13, IL3RA, IL27RA)* and cell type markers (*CD14*, *NKG7*, *GZMK*, *CD8B*). Taken together, our approach to identify DORCs uncovers genes under extensive chromatin control, likely a result of immune cells requiring exquisite control of transcription at these genes, thus reflecting key hubs of immunological response.

### Stimulated cells are characterized by early changes in the chromatin accessibility landscape that primes gene expression

Previously, we used multi-modal data to show that DORC accessibility foreshadows gene expression along developmental trajectories, and that this activity is predictive of cell state transitions^34^. To this end, we sought to use paired multi-omic data to ask whether cells prime for an immunological response through their chromatin accessibility states. Methods to deduce trajectory pseudotime often require the definition of a single root cell type. As we identified 18 discrete cell types, precluding the use of pseudotime, we sought to utilize an alternate approach to define trajectories. To do this, we computed a cell nearest-neighbor (NN) stimulation time per treatment which represents the weighted average of stimulus exposure time based on experimental treatment labels. Briefly, we take cell-nearest neighbors (*k*=50) for each cell, and compute the average neighborhood for control, 1h stimulated and 6h stimulated cells, assigning weights of 0, 1 and 2 respectively (**Fig 4A** and **Fig S5A**; see Methods). This continuous measure of time allows us to investigate the chromatin accessibility and gene expression dynamics along the stimulation trajectory.

**Figure 4.**
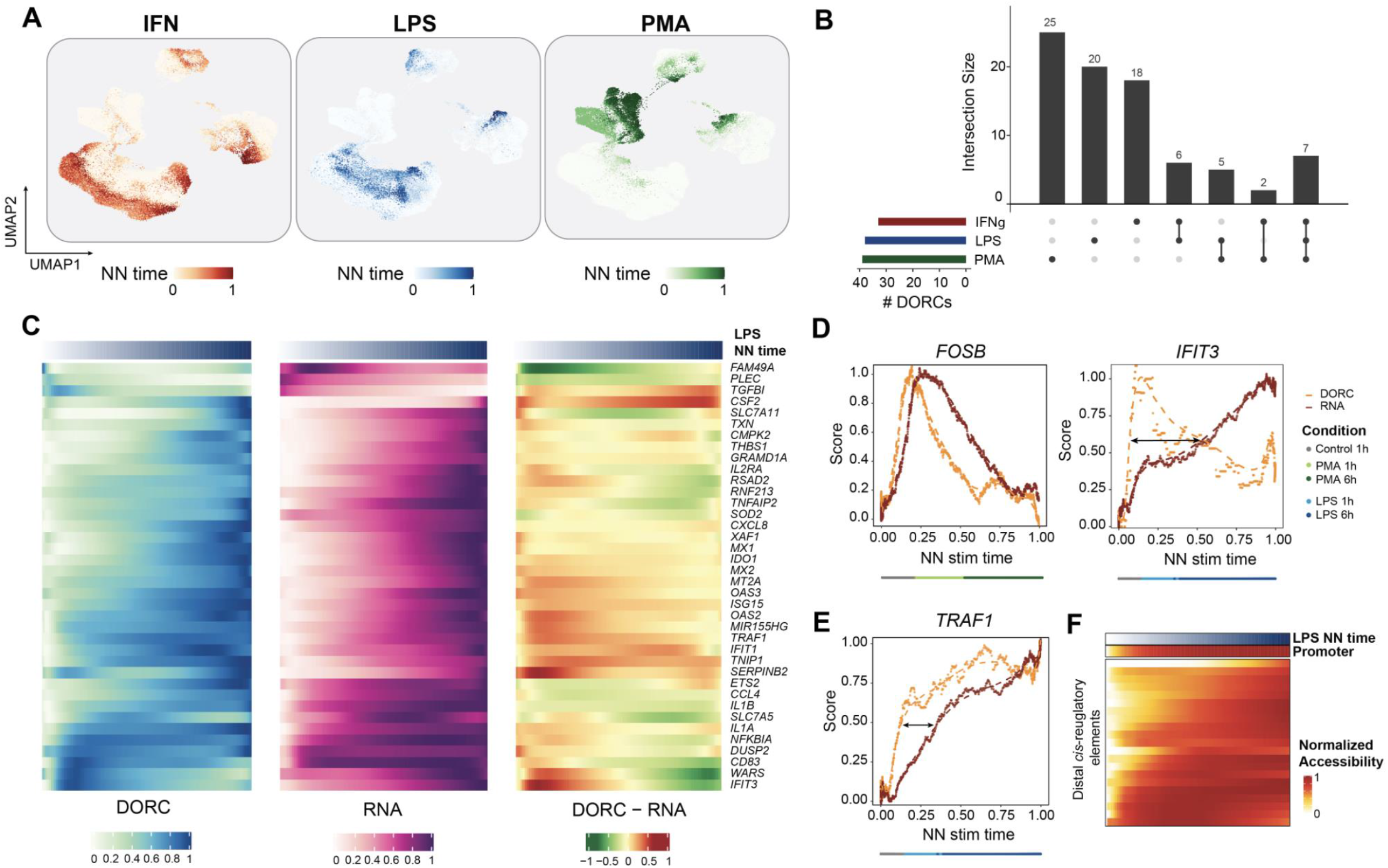
Chromatin and gene expression dynamics with respect to stimulus response time. **A.** UMAP of scATAC cells colored by estimated NN stimulation time per stimulus condition. **B.** UpSet plot highlighting overlap of monocyte-constrained DORC genes determined for the three different stimulus conditions. **C.** Heatmaps highlighting smoothed normalized DORC accessibility, RNA expression and residual (DORC - RNA) levels for DORC genes (*n*=38) identified to be associated with in LPS NN stimulation time in Control 1h and stimulated 1h/6h monocytes (*n*=1,776 cells). **D.** Chromatin (DORC) vs gene expression (RNA) dynamics of DORCs *FOSB* (left) and *IFIT3* (right) with respect to smoothed PMA and LPS NN stim time, respectively, for Control 1h and stimulated 1h/6h monocytes (*n*=2,002 cells for PMA+Control and *n*=2,601 cells for IFNγ+Control). Dotted line represents a loess fit to the values obtained from a sliding average of DORC accessibility or RNA expression levels (*n*=100 cells per sliding window bin). Color bar indicates the most frequent (mode) cell condition within each bin **E.** Same as in D, but for *TRAF1* with respect to LPS stimulated and Control 1h monocytes. **F.** Smoothed accessibility scores for individual *cis*-regulated elements correlated with *TRAF1* expression in control and LPS stimulated monocytes shown in Fig D, ordered by LPS NN stim time.

Using these stimulation time definitions, we sought to determine whether chromatin accessibility activates before gene expression, to thus “prime” or “foreshadow” immunological response. For this analysis, we chose to focus on the monocyte cellular population - as it is directly activated in response to our inflammatory factors, as described by prior literature and our observations with the Golgi Inhibitor (**Fig 3F-G**) - to assess chromatin and gene expression dynamics with respect to stimulation time. Restricting our peak-gene correlation approach strictly to control 1h, stimulation 1h and stimulation 6h monocytes, we identified a set of DORC genes associated with LPS (*n*=38 genes), IFNγ (*n*=33 genes), or PMA (*n*=39 genes) stimulation of monocytes. These DORCs include known expression markers induced upon stimulation in myeloid cells ^27^ Interestingly, we also found a small subset of these monocyte-specific DORCs are shared across multiple stimuli (**Fig 4B**). By averaging single-cell DORC accessibility and RNA levels in response to the NN stim time for each treatment (see Methods), we visualize the change of chromatin accessibility and gene expression along the control (0h) to 6h stimulation time axis (**Fig 4C** and **Fig S5C-D**).

Calculating the difference in chromatin versus RNA (residuals), we predominantly observed that chromatin accessibility in chromatin precedes that of expression (high residuals) at early time points. At later time points we found that residuals were low reflecting an accumulation of RNA following immune stimulation. These observations were stereotyped by the genes *FOSB* and *IFIT3* with LPS and PMA treatment, respectively (**Fig 4C-D** and **Fig S5E**). These changes occurred on relatively fast time-scales, for example priming of *FOSB* was an early event occurring early within the 60-minute time point. Notably, priming of DORC accessibility is constituted by individual *cis*-regulatory elements whereby some regulatory elements become accessible extraordinarily quickly (ie. the promoter) while others are slow to become accessible (some distal regulatory elements) along the stimulation time axis, as highlighted for LPS-responsive gene TNF receptor associated factor 1 (*TRAF1*) (**Fig 4E-F**). Conversely, we note a few exceptions including the PMA-responsive heat shock protein-encoding genes *HSP90AA1* and *HSPH1* which exhibit dominant early expression gain compared to the corresponding change in DORC accessibility (**Fig S5F**). Together, we demonstrate using computationally paired multi-omic data the ability to detect activation of chromatin accessibility prior to gene expression (“priming”) associated with stimulation-like cell states.

### A computational approach to identify candidate TF regulators of DORC activity

At the core of FigR, we developed a computational approach to define a gene regulatory network (GRN) of immunological response using multi-omic data. At this stage, FigR uses paired scATAC-seq and scRNA-seq data and specifically tests for the enrichment of TF motifs among predetermined *cis*-regulatory elements (i.e. DORCs), as well as the correlation of TF expression to the overall accessibility level for a given DORC gene (DORC score), to infer likely TF activators and repressors (**Fig 5A**). First, for a given DORC gene, FigR determines a pool of DORC *cis*-regulatory elements based on its DORC accessibility kNNs. This assumes that DORCs that are co-variable across the entire cell space are co-regulated by shared TFs. We then perform a statistical test for significance (*Z*-test) of TF motif enrichment using the frequency of motif matches across a reference database of TF motifs, relative to a background set of permuted peaks matched for GC content and global peak accessibility. Concomitantly, we compute the Spearman correlation coefficient between the TF RNA expression levels and the DORC accessibility score. Lastly, to determine activators and repressors, we combine significance estimates of relative motif enrichment (*Z*-test *P*) and RNA expression correlation (Z-test *P*) for a given DORC relative to all TFs, computing a signed probability score we term as a “regulation probability”, representing the intersection of both motif-enriched and RNA-correlated TFs. To enable the discovery of new regulators using this approach, we curated an expanded set of unique human (*n*=1,143) and mouse (*n*=895) TF binding sequence motifs, which extends upon a previously established database^37^ (see Methods).

**Figure 5.**
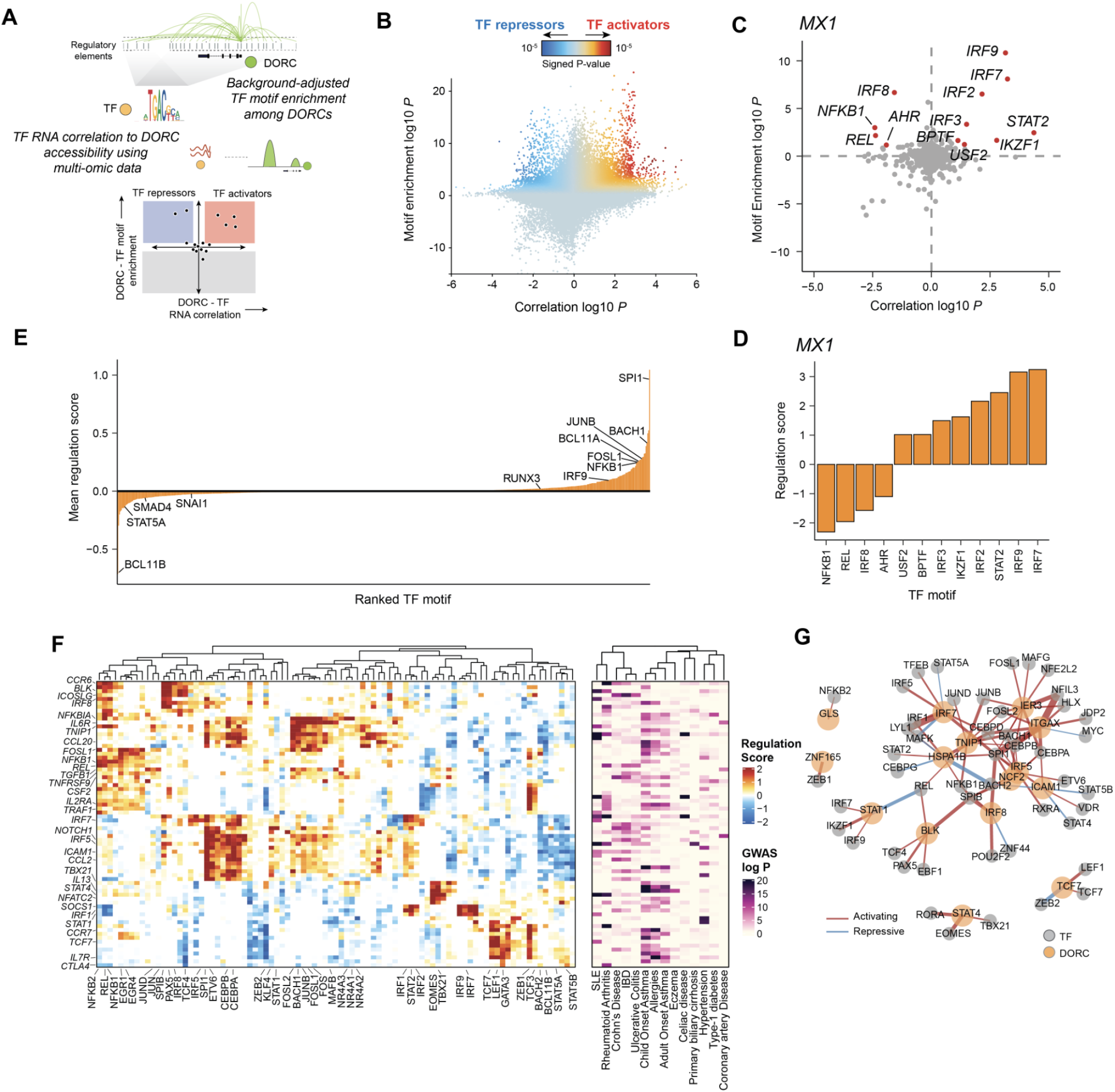
Design and application of the functional inference of gene regulation (FigR) workflow to identify TF modulators of immune response DORCs. **A.** Schematic describing FigR workflow. **B.** Scatter plot showing all DORC to TF associations, colored by the signed regulation score. **C.** Candidate TF regulators of *MX1*. Highlighted points are TFs with abs(regulation score) ≥ 1 (-log10 scale), with all other TFs shown in gray. **D.** Regulation scores (signed, -log10 scale) shown for highlighted TFs in C. **E.** Mean regulation score (signed, - log10 scale) across all DORCs (*n*=1,128) per TF (*n*=870), highlighting select TF activators (right-skewed) versus TF repressors (left-skewed). **F.** Heatmap of DORC regulation scores (left) for all significant TF-DORC enrichments for DORCs implicating GWAS variants (Absolute regulation score ≥ 1.5; *n*=89 TFs and *n*=73 DORCs). The corresponding min GWAS *P* (right; -log10 scale) for each DORC across all diseases considered is also shown. **G.** TF-DORC network visualization for SLE GWAS SNP-implicated DORCs (orange nodes), and their associated TFs (gray nodes) from F. Edges are scaled and colored by the signed regulation score.

To demonstrate the utility of FigR, we apply it to the paired stimulation scATAC-seq and scRNA-seq data to reveal key regulators of stimulus-response. To do this, we begin by testing all stimulus-responsive DORC genes (*n*=1,128) and reference TF motifs (**Fig 5B**). Filtering TF-DORC associations using a regulation score threshold (-log10 scale abs(regulation score) *≥* 1), we can then query putative TF regulators for a given DORC (**Fig 5C-D** and **Fig S6A-B**) as well as sets of DORCs that are potentially driven by a specific TF (**Fig S6C**). For example, FigR identifies known activators of *MX1*, including the IRF family of TFs *IRF3*, *IRF7, IRF9*, and *STAT2*, all belonging to the IFN signaling pathway^38^. We can further distinguish between TF activators versus TF repressors based on both their mean regulation score across all DORCs (**Fig 5E**), or by the fraction of positively and negatively regulated DORCs (**Fig S6D**). For example, we see *SPI1 (PU.1), BACH1,* and *BCL11A* as top transcriptional activators and *BCL11B* as a top transcriptional repressor (**Fig 5E** and **Fig S6D**). To demonstrate the broad generalizability of this approach, we applied it to multi-modal single cell data, using previously generated SHARE-seq data derived from murine skin tissue^34^ (**Fig S6E**). In doing so, FigR recovered TF regulators of DORCs which we previously found to associate with hair follicle differentiation. This includes activators *Lef1*, *Hoxc13,* and *Grhl1*, and repressors *Tcf12* and *Pou2f3*. Additionally, we determined activator *Dlx3*^39^, and repressors *Zeb1* and *Barx2*^40^ as top regulators (**Fig S6F-H**). Thus, we show that FigR can exploit both computationally or experimentally-paired multimodal data to derive GRNs using empirically-derived peak-to-gene and TF-to-peak motif associations to arrive at candidate TF regulators.

We next looked to see if the inferred stimulus response GRN from FigR may be used to reveal the regulatory mechanisms underlying disease associated genetic variants and their non-coding regulatory elements. To uncover disease associated cell states, we scored single cells for accessibility associated with GWAS SNP-overlapping peaks (GWAS *P* < 10^-7^; **Fig S6I**). We observed stimulus- and cell type-specific enrichment of chromatin accessibility for different inflammatory diseases tested (**Fig S6J**); validating prior work^18, 41^ showing that immunological stimulation uncovers regulatory elements enriched for disease GWAS variants. For example, we observed elevated enrichment of GWAS-associated accessibility in LPS and IFNγ-stimulated B lymphocyte and monocyte cells for systemic lupus erythematosus (SLE), and IFNγ- and PMA-stimulated CD4/CD8 lymphocytes for allergies (**Fig S6K**). Altogether, we find that our immunological stimulations uncover cell states, and their corresponding chromatin accessibility profiles, relevant to autoimmunity and associated genetic variation.

Next we reasoned that our GRN-based analysis may identify relevant mechanisms of disease-associated genetic variation. For example, the regulator NFkB is known to function across cell types to promote inflammatory gene expression^42^. Indeed, we found NFkB to drive activity of a large fraction of GWAS variant-associated DORCs (**Fig 5F**). Extending this analysis to all DORCs (*n*=77), we uncovered 89 putative TF drivers (abs(regulation score) *≥* 1.5) revealing a combination of lineage-determining as well as stimulus-responsive TFs spanning one or more diseases (**Fig 5F**). Closer inspection of the subset of SLE-specific DORCs (*n*=15 DORCs; *n*=48 associated TFs) revealed key regulatory associations, including previously determined SLE genes: *BLK, IRF5, IRF8 and NCF2* (**Fig 5G**). Taken together, our FigR approach can prioritize DORCs and their putative TF regulators to dissect the regulatory programs implicated among diverse autoimmune diseases. We include the inferred GRN to be interactively visualized through an R Shiny application (https://buenrostrolab.shinyapps.io/stimFigR/).

## Discussion

Here, we have generated a regulatory atlas of immunological stimulation in human blood. This effort was enabled both by high-coverage single-cell data, and the development of a new computational framework supporting multi-omic data integration, *cis*-regulatory analyses and the construction of an enhancer-aware GRN based on single cell profiles. In this effort, we overcame 3 key challenges: i) we implemented an approach to better computationally pair single cells, ii) we associate distal *cis*-regulatory peaks to target genes, and iii) we associate TFs to target genes. Importantly, the capability of FigR to synthesize GRNs using independently or concomitantly generated single-cell ATAC/RNA data will broadly enable GRN analyses across the broad range of scATAC-seq and related multi-omic technologies. Unlike prior methods that use co-expression or static measures of co-accessibility^21, 22^, GRN construction using FigR leverages both chromatin and RNA dynamics, through correlation of these features across single cells, providing a means to identify gene-regulatory relationships spanning cell states. To do this we utilize an empirical statistical approach to compute the probability of a TF-gene interaction, avoiding the use of heavily parameterized machine-learning approaches. Importantly, we also show the statistical tools described in FigR (peak-gene and TF-gene) are generalizable and can be applied to true multi-modal datasets assaying chromatin accessibility and gene expression from the same cell. However, we note that a limitation with this approach (and similar methods utilizing single-cell data) is that one can only determine regulatory relationships if they are variable across single-cells - constraining GRN models to observed changes across cell states, and precluding the analysis of “housekeeping” regulators.

We find that DORCs closely correspond to super enhancers (78%), and find that GWAS variants are enriched within DORCs that respond to immunological stimuli. Thus, defining peak-gene interactions and DORCs provides a useful platform to annotate the function of non-coding genetic variants corresponding to autoimmune and inflammatory conditions. Prior epigenomic studies have extensively utilized bulk analysis of histone modifications, chromatin accessibility and genome topology to annotate the function of disease-associated non-coding genetic variation^18, 28, 36, 43, 44^. Advancing beyond these prior studies, our single-cell multi-omic GRN approach provides a framework for associating key disease-associated loci to their target genes and regulating TFs. Further, a single-cell multi-omic approach enables the analysis of variant-associated DORCs and their putative TF drivers at single-cell resolution, enabling the discovery of rare cells harboring primed chromatin accessibility states at these disease-relevant genes.

Generally advancing the hypothesis that chromatin accessibility foreshadows gene expression (chromatin potential^34^), we find that chromatin accessibility precedes gene expression even with the extraordinarily fast gene expression dynamics associated with immunological stimulation. This, together with a large body of work^1, 45, 46^, upends the notion that chromatin change is “slow” or “stable” and instead paints a picture whereby chromatin structure is highly dynamic. Together we anticipate that our approach for defining GRNs will enable the elucidation of latent chromatin states that prime or inhibit cells from diverse environmentally-induced stimuli.

We anticipate future studies will further improve our capacity to predict gene-regulatory relationships using single-cell data. Specifically, multi-omic assays with even higher-coverage may further enable analyses that use TF footprinting information, as enabled by recent tools using ATAC-seq profiles^47^. These single-cell data-derived GRNs advance our ability to nominate essential regulators when employing emerging tools for single-cell functional genomics^48, 49^ and high-throughput perturbation strategies^50^. Overall, we envision a future of single-cell genomics that will shift towards studies of gene-regulatory processes advancing the predictive capability of cells undergoing fate transitions, as well as elucidating the latent/primed potential of cells prior to environmental stimuli, and their relevance in development and disease.

## Author contributions

V.K.K. lead all computational developments and analyses described in this work with contributions from Y.H., S.M., C.A.L. and Z.D.B. V.K.K and A.E. developed the online interactive resources. F.M.D., J.G.C. and A.S.K. generated the data with supervision by J.D.B and R.L. V.K.K and J.D.B. wrote the manuscript with input from all authors.

## Competing interests

J.D.B. holds patents related to ATAC-seq and scATAC-seq and serves on the Scientific Advisory Board of CAMP4 Therapeutics, seqWell, and CelSee. J.G.C., Z.D.B., A.S.K. and R.L. are employees of Bio-Rad.

## Methods

### Human peripheral blood mononuclear cells

Cryopreserved human peripheral blood mononuclear cells (PBMCs) and isolated peripheral blood CD4+, CD8+, CD14+, CD19+ and CD56+ cells were purchased from AllCells (see Table S1 for catalog numbers and donor information). Cells were quickly thawed in a 37°C water bath, rinsed with culture medium (Iscove’s Modified Dulbecco’s Medium (IMDM) (ATCC) supplemented with 10% FBS and 1% Pen/Strep) and then treated with 0.2 U/μL DNase I (Thermo Fisher Scientific) in 10mL of culture medium at 37°C for 30 min. After DNase I treatment, cells were washed with medium once and then twice with ice cold 1x PBS (Gibco) + 0.1% BSA (MilliporeSigma). Cells were then filtered with a 35 μm cell strainer (Corning) and cell viability and concentration were measured with trypan blue on the TC20 Automated Cell Counter (Bio-Rad). Cell viability was greater than 80% for all samples.

### Human PBMCs stimulations

PBMCs were quickly thawed in a 37°C water bath, rinsed with culture medium (RPMI 1640 medium supplemented with 15% FBS and 1% Pen/Strep) and then treated with 0.2 U/μL DNase I in 10mL of culture medium at 37°C for 30 min. After DNase I treatment, cells were washed with medium once, filtered with a 35 μm cell strainer and cell viability and concentration were measured with trypan blue on the TC20 Automated Cell Counter. Cell viability was greater than 90% for all samples. Cells were plated at a concentration of 1 x 10^6^ cell/mL, rested at 37°C and 5% CO_2_ for 1 h and then treated with the specified concentrations of the following stimulants (or DMSO as a control) for either 1h or 6h:

1. 20 ng/mL Lipopolysaccharide (LPS) (tlrl-3pelps, Invivogen),
2. 50 ng/mL Phorbol 12-myristate 13-acetate (PMA) (P8139, MilliporeSigma) + 250 ng/mL Ionomycin calcium salt (I0634, MilliporeSigma),
3. 20 ng/mL Interferon gamma (IFN-γ) (RP1077, Cell Applications)

For the “Golgi Inhibitor” experiments, cells were incubated for 6 h with GolgiPlug (555029, BD Biosciences) at a 1:1000 dilution plus stimulants at the concentrations indicated above (or GolgiPlug only as a control).

After stimulation, cells were washed twice with ice cold 1x PBS + 0.1% BSA and cell viability and concentration were measured with trypan blue on the TC20 Automated Cell Counter. dscATAC-seq experimental methods

### scATAC-seq experimental methods

#### Cell lysis and tagmentation

For a detailed description of tagmentation protocols and buffer formulations refer to the SureCell ATAC-Seq Library Prep Kit User Guide (17004620, Bio-Rad). Harvested cells and tagmentation related buffers were chilled on ice. Lysis was performed simultaneously with tagmentation. Washed and pelleted cells were resuspended in Whole Cell Tagmentation Mix containing 0.1% Tween-20, 0.01% Digitonin, 1x PBS supplemented with 0.1% BSA, ATAC Tagmentation Buffer and ATAC Tagmentation Enzyme (ATAC Tagmentation Buffer and Enzyme are both included in the SureCell ATAC-Seq Library Prep Kit (17004620, Bio-Rad)). Cells were mixed and agitated on a ThermoMixer (5382000023, Eppendorf) for 30 min at 37°C. Tagmented cells were kept on ice prior to encapsulation.

#### Droplet library preparation and sequencing

For a detailed protocol and complete formulations, refer to the SureCell ATAC-Seq Library Prep Kit User Guide (17004620, Bio-Rad). Tagmented cells were loaded onto a ddSEQ Single-Cell Isolator (12004336, Bio-Rad). Single-cell ATAC-seq libraries were prepared using the SureCell ATAC-Seq Library Prep Kit (17004620, Bio-Rad) and SureCell ddSEQ Index Kit (12009360, Bio-Rad). Bead barcoding and sample indexing were performed in a C1000 Touch™ Thermal cycler with a 96-Deep Well Reaction Module (1851197, Bio-Rad): 37°C for 30 min, 85°C for 10 min, 72°C for 5 min, 98°C for 30 secs, 8 cycles of 98°C for 10 secs, 55°C for 30 secs, and 72°C for 60 sec, and a single 72°C extension for 5 min to finish. Emulsions were broken and products cleaned up using Ampure XP beads (A63880, Beckman Coulter). Barcoded amplicons were further amplified using a C1000 Touch™ Thermal cycler with a 96-Deep Well Reaction Module: 98°C for 30 secs, 6-9 cycles (cycle number depending on the cell input, Section 4 Table 3 of the User Guide) of 98°C for 10 secs, 55°C for 30 sec, and 72°C for 60 sec, and a single 72°C extension for 5 min to finish. PCR products were purified using Ampure XP beads and quantified on an Agilent Bioanalyzer (G2939BA, Agilent) using the High-Sensitivity DNA kit (5067-4626, Agilent). Libraries were loaded at 1.5 pM on a NextSeq 550 (SY-415-1002, Illumina) using the NextSeq High Output Kit (150 cycles; 20024907, Illumina) and sequencing was performed using the following read protocol: Read 1 118 cycles, i7 index read 8 cycles, and Read 2 40 cycles. A custom sequencing primer is required for Read 1 (16005986, Bio-Rad; included in the kit).

### scRNA-seq experimental methods

Single-cell RNA-seq (scRNA-seq) data for LPS, PMA or IFNγ-stimulated cells, and isolate (bead-enriched) PBMCs comprising CD19^+^, CD4^+^ T-cells, CD8^+^ T-cells, CD56^+^ Natural Killer (NK) cells and CD14^+^ monocytes were generated using the SureCell WTA 3′ Library Prep Kit for the ddSEQ System (20014280, Illumina) with the following modifications. A higher concentration of beads was used to obtain 1,000-2,000 single-cells per emulsion, whilst minimizing the number of droplets with multiple beads to < 10%. Furthermore, Bst 2.0 WarmStart (M0538S, NEB) was added to the droplet mix to perform temperature activated second strand synthesis in droplets.

### scATAC-seq analysis workflow

#### Raw read processing, demultiplexing and alignment

Per-read bead barcodes were parsed and trimmed using UMI-TOOLs (https://github.com/CGATOxford/UMI-tools)^51^, and the remaining read fragments were aligned using BWA (https://github.com/lh3/bwa) on the Illumina BaseSpace online application. Constitutive elements of the bead barcodes were assigned to the closest known sequence allowing for up to 1 mismatch per 6-mer or 7-mer (mean >99% parsing efficiency across experiments). All sequence libraries were aligned to the hg19 reference genome. We then used bead-based ATAC-seq data processing (BAP, v0.6.4) (https://github.com/caleblareau/bap)^13^ to help identify systematic biases (i.e. reads aligning to an inordinately large number of barcodes), barcode-aware deduplication of reads, and to perform merging of multiple bead barcode instances associated with the same cell (barcode merging is necessary due to the nature of the Bio-Rad SureCell scATAC-seq procedure used in this study, which enables multiple beads per droplet). For a detailed description of the bead barcode merging strategy see^13^. We ran BAP using a single input alignment (.bam) file for a given experiment with a bead barcode identifier indicated by the SAM tag “DB”, and default parameters.

#### Chromatin accessibility peak calling

Genome-wide chromatin accessibility peaks were called using MACS v2 (MACS2)^52^ on the merged aligned scATAC-seq reads per treatment condition, generating a list of peak summit calls per condition. As previously described, summits were then ranked per condition based on their FDR score (from MACS2), and the summit scores rank-normalized such that the normalized summit scores rendered are comparable across conditions ^11^. Peak summits were then padded by 400 bases on either end (generating 801 bp windows), and overlapping peak windows filtered iteratively such that windows with higher scores were retained at each step. This resulted in a filtered list of disjoint peaks (*n*=219,136), which were finally resized to 301 bp (i.e. ± 150 bp from each peak summit) and used for all downstream analyses.

#### scATAC-seq counts generation and QC

Single cell counts for reads in peaks were generated by intersecting the peak window regions (see previous section) with aligned fragments. First, we offset the start and end coordinates of the aligned fragments to identify Tn5 cut sites by +4 or -5 bp for fragments aligning to the positive or negative strand, respectively. These are then intersected with peak window regions using the findOverlaps function in R, and the total number of unique fragment cut sites overlapping a given peak window tallied for each unique cell barcode detected in the data, producing a matrix of single cell chromatin accessibility counts in peaks (rows) by cells (columns). Only cells with fraction of total reads in peaks (FRIP) ≥ 0.5, a minimum of 2,000 unique nuclear fragments (UNFs), and a sequencing library duplication rate ≥ 0.15 were retained. Cell barcode doublets were inferred and filtered out using ArchR^31^. This resulted in a total of *n*=67,581 and *n*=17,920 cells, for the stimulated and isolate PBMC cells, respectively.

#### TF motif scoring

Single cell accessibility scores for TF motifs were computed using chromVAR^37^, as also previously described^12, 13^. For TF motif accessibility scores, the peak by TF motif overlap annotation matrix was generated using a list of human TF motif PWMs (*n*=870) from the chromVARmotifs package in R (https://github.com/GreenleafLab/chromVARmotifs), and used along with the scATAC-seq reads in peaks matrix to generate accessibility Z-scores for across all scATAC-seq stimulated cells passing filter.

#### Gene TSS scoring

Single cell gene activity scores were generated using scATAC-seq data based on an exponential decay weighted sum of fragment counts around a given gene TSS using a previously described approach^53^ for all scATAC-seq cells passing filter using the hg19 reference for gene TSSs. Raw gene scores were then normalized by dividing by the mean gene score per cell.

#### scATAC-seq cell clustering, visualization and annotation

Single cell clustering of ATAC-seq data was performed using the ArchR framework^31^. First, the accessibility counts in a tiled window matrix was determined using default parameters. ArchR’s iterative LSI dimensionality reduction was performed for *n*=30 components and *n*=2 iterations, taking the top variable 50000 peaks and evaluating resolutions 0.1 to 0.4, sampling 20,000 cells. Cells were projected in 2D space using uniform manifold approximation and projection (UMAP), based on the top 30 LSI components with the addUMAP function (nNeighbors=50, metric=”cosine”, min.dist=0.5). These steps were applied independently for both stimulation cell scATAC-seq cells and PBMC isolate cell scATAC-seq cells passing filters. Annotation of stimulation scATAC-seq cells was obtained using the corresponding annotation of paired scRNA-seq cells (see sections ‘scRNA-seq cell cell clustering, visualization and annotation’ and ‘scRNA-seq and scATAC-seq OptMatch pairing’ below for more details). For isolate PBMC scATAC-seq cell clustering, the same LSI and UMAP parameters were used to obtain 2D clustering of cells based on peak accessibility.

#### GWAS variant enrichment analyses

Summary statistics for 12 of the 14 GWAS traits were downloaded from sources as previously described^54^. The remaining traits were downloaded from the SAIGE resource (Adult/Child onset asthma)^55^ or the EAGLE consortium (Eczema)^56^. Raw summary statistics were then reformatted uniformly for downstream analyses and processing, including a per-SNP association p-value threshold of *P* < 10^-7^ for the list of final variants considered for peak-SNP overlaps. For each trait considered, filtered variant loci were intersected with peaks using the findOverlaps R function, to generate a peak by variant binary overlap matrix. This was then multiplied by a variant by trait binary annotation matrix to yield a peak by trait annotation matrix. This resulting annotation matrix was used, along with scATAC-seq reads in peak counts, as input to chromVAR to generate single cell trait *Z*-scores based on the relative enrichment of ATAC-seq counts within these trait-associated peaks (used for UMAP visualizations in Fig S6J). For aggregate SNP scores, single cell *Z*-scores were converted to one-tailed p-values using a *Z*-test, and the resulting p-values combined using the Fisher method^57^, per condition and cell type, and used for heatmap visualizations (related to Figure S6K).

### scRNA-seq analysis workflow

#### Raw read processing, demultiplexing and alignment

The library preparation for scRNA-seq experiments configures the reads such that read 1 contains a cell barcode and UMI and read 2 the cDNA generated from the transcript. Cell barcodes and UMIs were parsed from read 1 and written into the read name of the corresponding read in the read 2 fastq. All read 2 files with valid cell barcodes were aligned using STAR (v2.5.2b) to hg19 (UCSC; PAR masked) reference genome. Reads that aligned to abundant features (chrM, rRNA, and sncRNA) were filtered from the analysis.

#### scRNA-seq counts generation and QC

Transcript counts per barcode were then generated by counting the number of unique genic UMIs for each read with a minimum mapping quality of 12 that aligned unambiguously to an annotated exon in the RefSeq annotation of hg19. The distribution of unique genic UMI per barcode was then filtered to separate barcodes present in droplets with cells from barcodes present in cell-free droplets. First, a background filtration step was performed to remove barcodes that arose from sequencing errors and empty droplets by computing a background threshold. The background threshold was computed to filter barcodes that arose from sequencing errors and empty droplets by performing a kernel density estimate on the log10 transformed genic UMIs per barcode distribution wherein the largest peak is assumed to be from cell-free droplets. The number of UMIs corresponding to this peak was deemed the “background level”. The half-height of the background peak was calculated by measuring the distance from the top of the background peak to the point on the right where the density dropped to 50% of the peak. The standard deviation of the background peak was then estimated by dividing the half-width by 1.17 under the assumption of the background peak being a normal distribution. Finally, the background threshold was calculated as the background level + 5 * the standard deviation of the background peak. All barcodes below this value were filtered from the analysis.

After background filtration, the remaining barcodes in the genic UMI count distribution were subjected to a “knee calling” algorithm wherein inflection points in kernel density estimate of the log10 transformed UMI count distribution were identified. The leftmost inflection point (= higher genic UMI count) was used to determine the final cell count.

Gene-mapped counts were then loaded into R as a Seurat object^29^ and used for downstream analysis. Genes with at least 1 UMI across cells were retained, and cells with number of unique feature counts > 200 and < 5000 were initially retained. Normalization and scaling of RNA gene expression levels was performed using the SCTransform function. scRNA cell barcode doublets were inferred using DoubletFinder^58^ and removed.

#### scRNA-seq cell clustering, visualization and annotation

For the stimulation scRNA-seq cell clustering (shown in Fig 1), PCA was first run on the normalized scRNA-seq counts using the runPCA function in Seurat. The first 30 PCs were then used to run UMAP for single cell 2D projection using Seurat’s RunUMAP function. For stim-corrected clustering of scRNA-seq cells (Fig S2, used for cell type annotation), we followed Seurat’s workflow for integrating batches using canonical correlation analysis (CCA), where we treated each condition (e.g. Control 1h or LPS 6h) as a batch, following the integration protocol steps for finding cell integration anchors with default settings (https://satijalab.org/seurat/archive/v3.1/immune_alignment.html). The corresponding batch-aligned integrated data was scaled, and PCA dimensionality reduction was run. UMAP was used for the final cell projection (top 30 PCs, min.dist=0.5), and a cell kNN graph was determined using the FindNeighbors function in Seurat (*k*=10 cell neighbors). Cells were then grouped into clusters using the FindClusters Seurat function (resolution=0.8; SLM algorithm), and cluster and cell annotations manually assigned by visualizing the mean and percent expression of cell identity markers within cell clusters (Fig S2). Broader annotations (e.g. monocytes) were determined by merging finer cell groupings (e.g. CD14 and CD16 monocytes). For isolate PBMC scRNA-seq cell clustering, the same PCA and UMAP parameters were used to obtain 2D clustering of cells.

### scRNA-seq and scATAC-seq OptMatch pairing

Computational pairing of scATAC and scRNA cells was performed either per treatment condition (stimulation data) or across all cells (PBMC isolates) using an approach we refer to throughout as “OptMatch”. First, the union of the top 5,000 variable genes based on genescore (ATAC) and gene expression (RNA) was taken across all cells, determined using Seurat’s FindVariableFeatures function on normalized scATAC genescores and normalized scRNA gene expression. These features were then used to perform a canonical correlation analysis (CCA) using the RunCCA function. The L2-normalized CCA components (*n*=30) were then visualized using UMAP to highlight co-embedding of the two assays for the same cellular context (Fig S3). This was done for both the PBMC isolate data (*n*=17,920 ATAC, *n*=8,089 RNA cells), as well as the stimulation data (*n*=67,581 ATAC, *n*= 23,754 RNA cells).

Next, to (globally) balance ATAC and RNA cell numbers, we first randomly divide the larger (in our case, ATAC) dataset into chunks of cells size equal to the original number of cells in the smaller (in our case, RNA) dataset, re-sampling cells from the RNA cell pool to match the remainder (unsampled) ATAC cells for the final smaller cell chunk. Then, for each generated cell chunk having the same number of sampled ATAC and RNA cells, we rederive a 5D UMAP cell embedding based on the CCA components (1 to 20; *k*=30 cell neighbors) using the uwot R package, and determine for each cell an undirected *k*-nearest neighbor (kNN) graph (*k*=5 cell neighbors) based on the 5 UMAP embedding dimensions. Using this neighbor graph, we determine the shortest path distance (geodesic distance) between all cells using the shortest.paths function in the igraph R package, using which we divide the cell chunk into connected subgraphs (subclusters with finite non-zero geodesic distance) using the clusters function in the igraph package, only retaining subgraphs of size 50 cells or more. For each subgraph, we then deal with assay cell type imbalance by matching the number of local ATAC/RNA cells through random sampling of the smaller to the larger dataset, without replacement, yielding equal cells for ATAC and RNA in the subgraph.

To greatly reduce computational complexity of optimal matching (traveling salesman problem), we implement a sparse-kNN matching approach by only pairing ATAC-RNA cells that are within a geodesic distance kNN range (k*_g_*) from each other in the subgraph, where the threshold k*_g_* is set as:

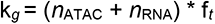

where

f*_t_ =* Fraction of total cells in the subgraph to consider as geodesic kNN upper-bound (set to 0.1)

*n*_ATAC_ = # ATAC cells in subgraph

*n*_RNA_ = # RNA cells in subgraph

Finally, using the geodesic distance as a cost function, we determine the optimal pairing within the established geodesic ATAC - RNA kNNs subgraph cell space using the fullmatch function in the optmatch R package (https://github.com/markmfredrickson/optmatch)^32^, setting the following non-default parameters: tol: 0.0001, max_multimatch=5.

The overall performance of the OptMatch approach described above for pairing single cells across ATAC/RNA datasets was assessed using previously published scATAC-seq data ^13^ from cells sorted for CD19^+^, CD4^+^ T-cells, CD8^+^ T-cells, CD56^+^ Natural Killer (NK) cells and CD14^+^ monocytes, and newly generated scRNA-seq data from the same cell pool for each enriched cell population (see scRNA-seq experimental methods). For comparison, we also determined a “greedy” assignment of cell pairs, for which we assigned each cell in the ATAC dataset to the cell with the highest similarity score in the RNA dataset (maximum Pearson correlation based on the first 20 CCA components). Overall performance between the two pairing modes was determined based on the percentage accuracy based on matching concordant cell types across assays for each sorting experiment (e.g. how often a CD19^+^ cell in the ATAC dataset paired with a CD19^+^ cell in the RNA dataset), the frequency of ‘multi-matches’ (multiple RNA cells pairing to a single ATAC cell), and the final percentage of paired cells in both ATAC and RNA datasets. Pairs were visualized by picking 300 ATAC-RNA pairs at random, highlighting the corresponding cells in CCA UMAP space.

### Aggregate ATAC and RNA profiles

Paired aggregate single cell peak (scATAC-seq) and gene (scRNA-seq) “pseudobulk” counts for different conditions and cell types (see Fig 3E) were obtained by summing the normalized scATAC-seq peak accessibility counts separately for promoter peaks (peak windows found within 1000 bp upstream and 300 bp downstream from each gene’s TSS, using the promoters function in the GRanges package) and distal peaks (peaks found outside defined promoter window), and by summing Seurat-normalized scRNA-seq gene counts across cells per condition and cell type. These pseudobulk counts were then quantile-normalized to adjust for differences in overall cell numbers across groupings.

### Peak-gene *cis*-regulatory correlation analysis

High density domains of regulatory chromatin (DORCs) were determined using scATAC-seq and scRNA-seq data for computationally-paired cells (see section above). Briefly, a 100 kb window was taken around the TSS of all hg19 RefSeq genes that were found to be expressed based on scRNASeq data. Next, peak-gene pairs where peak summits overlapped a given gene TSS window were determined (*n* = 155,831 peaks and 18,151 genes and a total of 343,640 gene-peak pairs). For each pair, the observed gene-peak correlation coefficient (Spearman ⍴) was determined by correlating the mean-centered scATAC-seq peak counts with the corresponding gene’s expression across all ATAC-RNA paired cells (*n*=62,219 cells). Permuted correlation coefficients for each gene-peak pair were calculated using background peaks matched for GC content and total chromatin accessibility levels across cells for each peak tested, determined using chromVAR (*n*=100 iterations). Finally, the significance of each gene-peak association was determined using a one-tailed *Z*-test computed from the observed and permuted coefficients. Only gene-peak associations that show positive correlations and were statistically significant (*Z*-test permutation *P* ≤ 0.05) were considered, and used to identify DORCs based on the number of significant peaks associated with each gene (DORCs = genes with n ≥ 7 associated peaks). Single cell DORC scores per gene were calculated as the sum of normalized scATAC-seq reads in peak counts (mean-centered) using the corresponding significantly correlated DORC-peaks for that gene, and smoothed for visualization based on *k*=30 cell kNNs derived using the scATAC-seq LSI components.

### DORC super-enhancer analysis

To determine overlap of stimulation DORCs and previously annotated super-enhancer regions, we used a previously annotated ^43^ list of genes associated with super-enhancers spanning different cellular contexts (*n*=86). We then determined and visualized the cumulative fraction of all stimulus DORCs (*n*=1,128) that overlap with each of the different super-enhancer linked gene lists.

### Differential DORC analyses

For differential testing of DORC accessibility scores or expression levels, we used normalized single cell DORC scores (paired scATAC-seq cells; *n*=62,219), or RNA expression (unpaired scRNA-seq cells; *n*=23,754) and performed differential testing using a Wilcoxon rank sum test per cell type (CD4/CD8 T, B, Monocyte, and NK), comparing each stimulus condition to its corresponding control condition (e.g. IFNγ 1h vs Control 1h for Monocytes) for all determined DORCs (*n*=1,128 genes). FDR was determined to adjust for multiple tests. For visualization, only the union of top 10 genes (ranked by nominal DORC ATAC Wilcoxon test *P*) per comparison were kept (*n*=53 DORCs), and a heatmap of the difference in mean single cell score (DORC accessibility or RNA expression) was used, showing the most significant change across any of the five cell types assessed for each DORC and condition, along with the corresponding cell type which reported the minimum *P* across any condition for each DORC.

### Cell nearest neighbor (NN) stimulation time calculation

Cell NN stimulation time for scATAC-seq cells was computed based on a weighted average of cell-nearest neighbor conditions. For each scATAC-seq cell in the paired stimulation data (*n*=62,219), we used the first 30 LSI components to derive a *k*=50 nearest neighbor (NN) graph as the cell’s nearest condition cells, leaving out the Golgi inhibitor treatment condition. Then, for each cell and it’s kNNs, we computed the mean stimulation time as the weighted average of its kNNs, using a weight of 0,1 and 2 for Control 1/6 hour, stimulation 1hr and stimulation 6hr time points respectively, done separately for each of the three stimulus conditions (LPS, IFN or PMA). The resulting estimates were then rescaled to fall between 0 and 1, and used for downstream analyses including fitting and visualization of DORC and RNA expression values to NN stimulation time.

### DORC accessibility and RNA expression dynamics

To visualize dynamics of DORC accessibility and gene expression along the NN stimulation time axis, we took scATAC-seq cells annotated as monocytes pertaining to control 1h, as well as stimulation 1h and stimulation 6h time conditions for LPS (*n*=1,776 cells), IFNγ (*n*=2,601 cells) and PMA (*n*=2,002 cells). Using a window size of *n*=100 cells, we then computed the rolling average DORC accessibility and gene expression value, which was then min-max normalized to the 1-99 percentile value, respectively. Additionally, we fit a loess smoothing function (loess alpha=0.1) using the normalized DORC/RNA values to the smoothed (rolling average) NN stimulation time, which was overlaid and visualized.

### Motif database

From cisBP, we curated position frequency matrices (PFMs) that represented a total of 113,635 human motifs and 107,308 mouse motifs. We filter motifs to a unique subset, one motif for each TF regulator, resulting in 1,143 unique human or 895 unique mouse TFs and motifs. To do this, we iterated through each unique TF to find all associated motifs from the high-quality motif list (as annotated by cisBP). For these associated high-quality motifs, we computed a similarity matrix using the Pearson correlation of the PFMs. To select the most representative motif for each TF regulator, we found the motif correlated with the most other motifs of the same TF at R > 0.9. If a TF was not represented in the high-quality list, we repeated the process using the medium- and low-quality databases for TF regulators. The final curated motif database contains 1,141 human and 890 mouse unique regulators and motifs, and can be incorporated with FigR and other PFM utilizing packages.

### FigR workflow

To associate TF regulators to target DORC gene activity, we deduced a metric that combines the relative enrichment of TF motifs among DORCs and the correlation of TF RNA expression with DORC accessibility. First, for each of our defined DORC genes, we determine a reference pool of expression-correlated chromatin accessibility peaks associated with its *k*-nearest neighbor DORC genes (default *k*=30). Then, for each DORC, we use its pooled peak set to perform an enrichment *Z*-test of the observed TF motif-to-peak match frequency with respect to each TF in our curated TF motif list described earlier (*n*=870), relative to the expected frequency based on matches to a permuted background peak set matched for GC content and overall accessibility (default *n*=50 permutations). We then correlate across all paired cells (*n*=62,219) the smoothed DORC accessibility score with the smoothed paired RNA expression levels of all tested TFs (smoothed using *k*=30 nearest cell neighbors based on first 30 LSI components), and use the standardized Spearman correlation levels to perform a *Z*-test to establish significance of correlation. Lastly, we combine the two significance levels (correlation and peak enrichment) and define a “regulation score” in log space as follows:

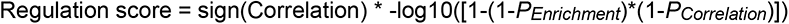

All regulation scores corresponding to negative TF enrichments (TF enrichment *Z*-score < 0) were set to 0.

For the stimulation GRN, putative regulators of DORCs were defined as TFs that have an absolute regulation score ≥ 1. SNP-DORC regulatory associations were visualized by taking the list of DORCs whose associated peaks overlap any disease GWAS variant (*n*=77 DORCs) and clustering DORC-TF associations with absolute regulation score ≥ 1.5 (*n*=73 DORCs and *n*=89 TFs). Network plots for a subset of DORCs (e.g. SLE-specific DORCs) were drawn for the associated filtered edge-node associations using the ggnet2 R package, and can be further visualized through the R Shiny App (https://buenrostrolab.shinyapps.io/stimFigR/). For the murine skin tissue GRN, previously published SHARE-seq data and the corresponding DORC calls were used^34^, along with the mouse cisBP TF motif database (*n*=797 motifs), with *k*=20 DORC kNNs for peak pooling.

## Data Availability

Analysis code can be found on GitHub (https://github.com/buenrostrolab/stimATAC_analyses_code). Additionally, gene regulatory networks and single cell profiles (cell metadata, DORC scores and paired RNA expression) for stimulation data can be interactively queried through our R Shiny app (https://buenrostrolab.shinyapps.io/stimFigR/). Newly curated TF motif PFM lists for human and mouse are available for download and use as R PFMList objects on github. Normalized aggregate scATAC-seq coverage profiles (BigWig) can be visualized for monocytes per condition (https://genome.ucsc.edu/s/vkartha/stimATAC_CD14_conditions) or for control 1h cells across paired cell type annotations (https://genome.ucsc.edu/s/vkartha/stimATAC_Control1h_cellTypes) using the UCSC genome browser.

## Acknowledgements

We thank members of the Buenrostro lab and Bio-Rad team for useful discussions and critical assessment of this work. J.D.B. and the Buenrostro lab acknowledge support from the Gene Regulation Observatory at the Broad Institute of MIT & Harvard, the Chan Zuckerberg Initiative, and the NIH New Innovator Award (DP2).

---

**Figure S1 (related to Figure 1).**
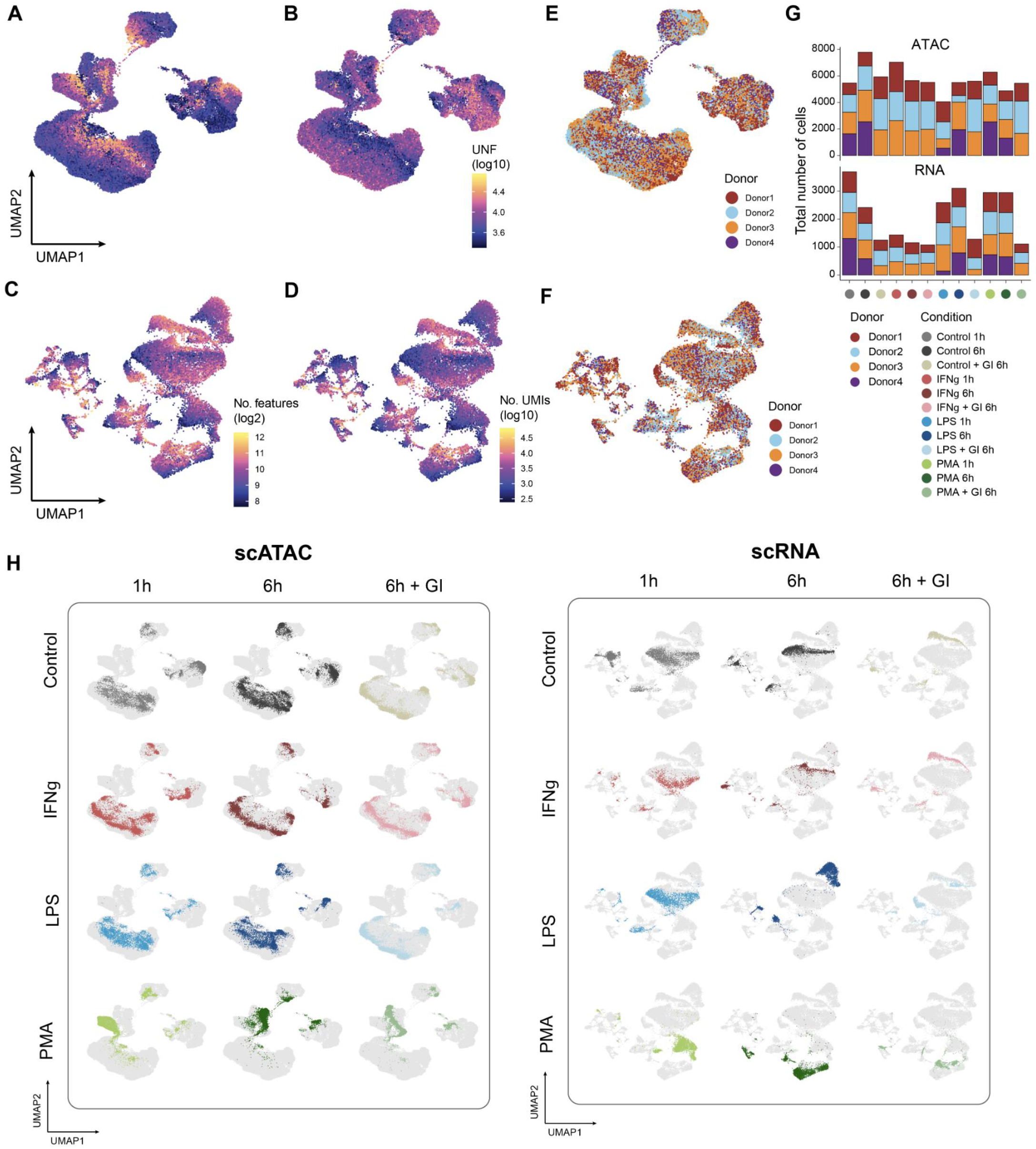
**A-B.** UMAP projection of scATAC-seq cells colored by fraction of reads in peaks (FRIP) (A) or total number of unique nuclear Tn5 insertion fragments (B). **C-D.** UMAP projection of scRNA-seq cells colored by total number of detected features (C) or total number of unique molecular identifiers (UMIs) per feature (D). **E-F.** UMAP projection of scATAC-seq (E) and scRNA-seq (F) stimulation data colored by Donor. **G.** Number of cells passing quality filtering for scATAC-seq and scRNA-seq stimulation data per donor per condition. **H.** UMAP of scATAC-seq (left) and scRNA-seq (right) cells profiled, with cells for each condition highlighted on the background of all cells.

**Figure S2 (related to Figure 1).**
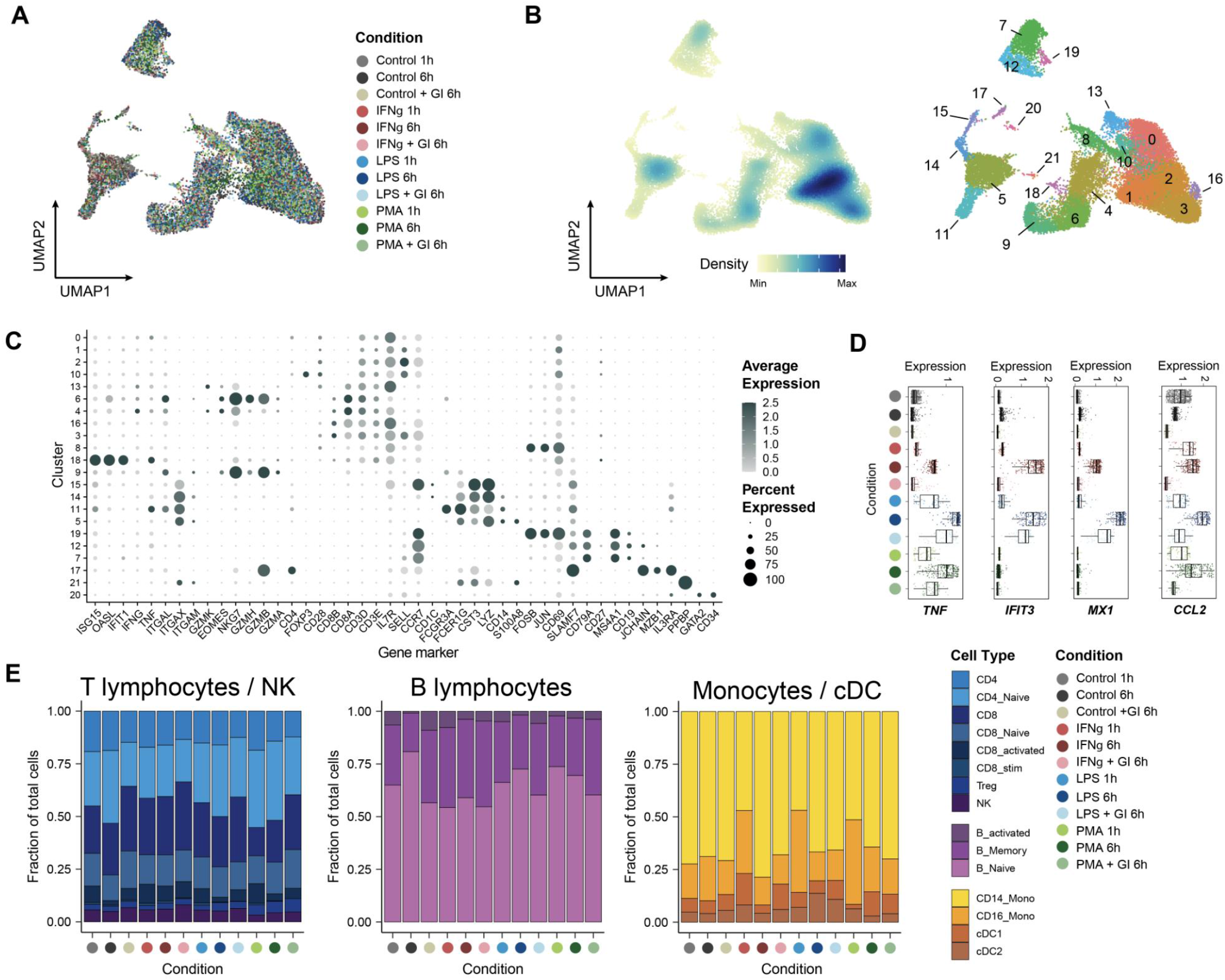
**A.** UMAP of scRNA-seq cells (aligned) after adjusting for treatment condition **B.** UMAP of aligned scRNA-seq cells in A, colored by density (left) or cell cluster based on Leiden clustering (right). **C.** Dotplot of gene expression markers highlighting cluster specific expression (used for scRNA-seq cell annotation) **D.** Smoothed RNA expression distribution in CD14 Monocytes across conditions for specific stimulus response genes **E.** Fraction of total scRNA-seq CD4/CD8/NK cells (left), B lymphocytes (middle) and Monocytes / cDCs (right) grouped per condition and cell type annotation.

**Figure S3 (related to Figure 2).**
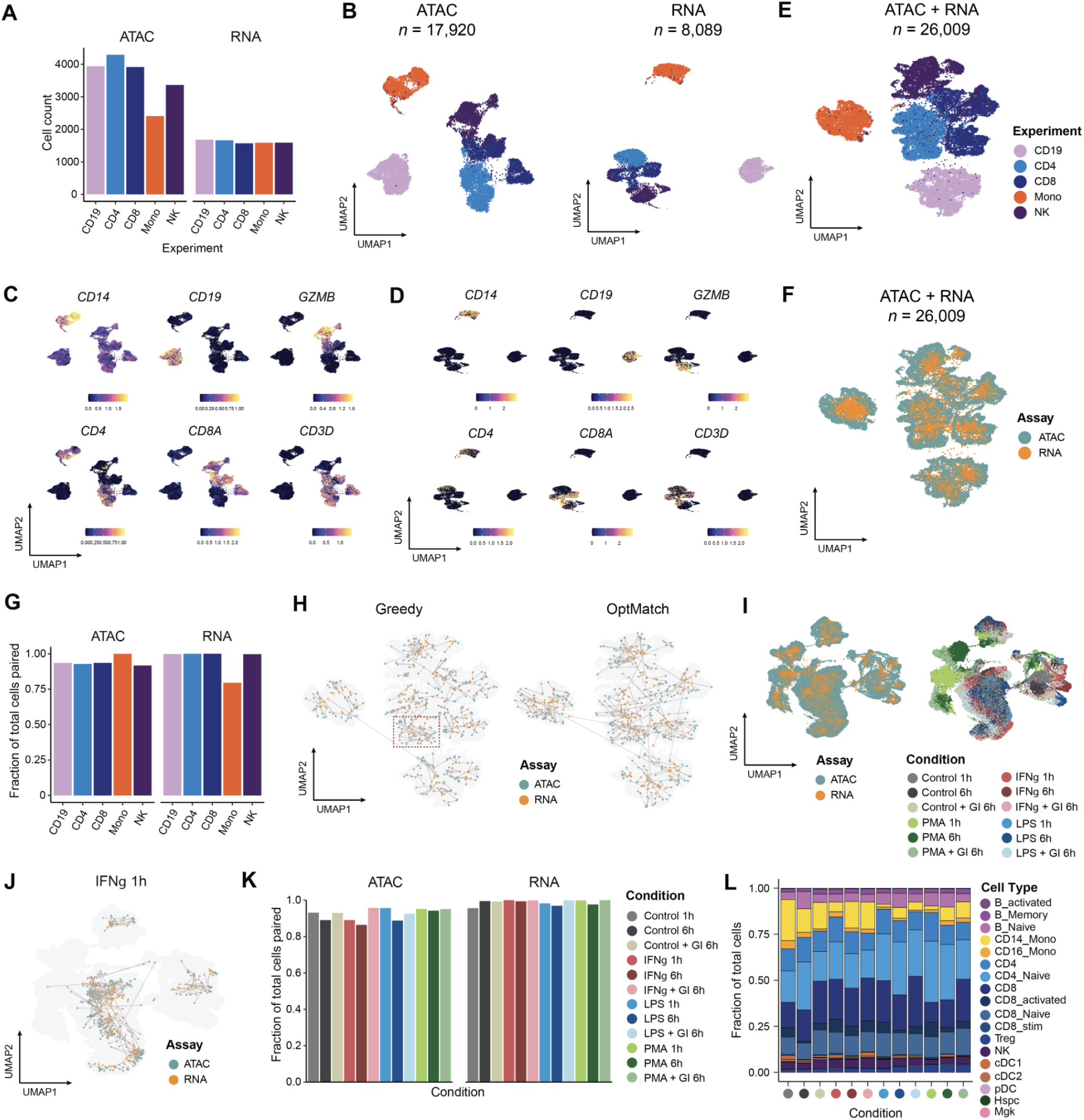
**A.** Total number of cells assayed and passing QC for scATAC-seq and scRNA-seq from bead enriched cells **B.** UMAP plots of bead enriched PBMCs showing single cell projections for scATAC (left) or scRNA cells (right), based on peak accessibility or gene expression, respectively. **C.** UMAPs of scATAC cells from B colored by smoothed gene activity scores of cell type marker genes **D.** UMAPs of scRNA cells from B colored by gene expression levels for cell type marker genes. **E.** UMAP of both scATAC and scRNA cells based on CCA co-embedding using union of top variable scATAC gene scores and top variable scRNA gene expression, with cells colored by enriched sub-population **F.** Same as in E, with cells colored by assay. **G.** Fraction of total scATAC and scRNA-seq cells paired using our OptMatch approach per isolate cell type assayed. **H.** CCA-based UMAP of cells from F, highlighting computational pairing (300 pairs shown at random) between scATAC-seq and scRNA-seq cells using a greedy approach (scRNA cell with maximum Pearson r for each scATAC cell; left) versus our OptMatch constrained pairing method (right). Red box highlights multiple RNA cells mapping to the same ATAC cell **I.** UMAP clustering based on CCA co-embedding of stimulation data cells colored by assay (left) or by stimulus condition (right). **J.** Computational pairing (300 pairs shown at random) between scATAC-seq and scRNA-seq cells for the IFN 1h stimulation condition. **K.** Fraction of total scATAC and scRNA-seq cells paired using our OptMatch approach per stimulus condition assayed. **L.** Distribution of scATAC cells (*n*=62,219) based on paired annotation obtained from pairing to scRNA-seq cells, per condition.

**Fig S4 (related to Figure 3).**
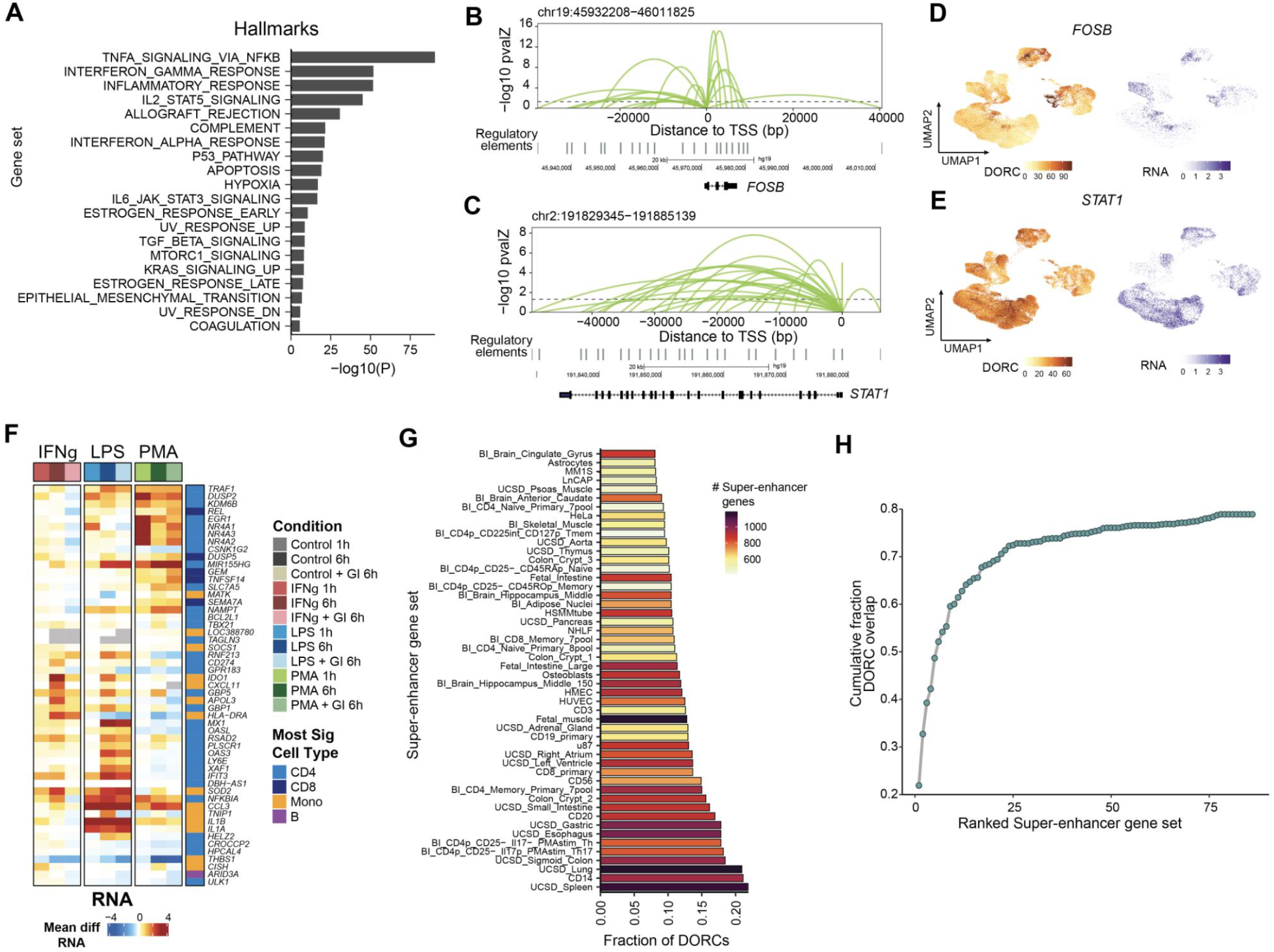
**A.** Gene set hyper-enrichment testing among DORCs for Hallmark gene sets (*n*=50). **B-C.** Loop plots highlighting significant gene-peak associations for *FOSB* (B) *and STAT1* (C). **D-E**. UMAPs of paired cells (*n*=62,219) highlighting scATAC DORC scores (left) or scRNA expression (right) for *FOSB* (D) and *STAT1* (E). **F.** Heatmap of the mean difference in single cell RNA expression for the union of the top 10 differential DORCs across conditions and cell types (*n*=53 genes; see Fig 3G). Cell type color bar represents the cell group having the most significant change across all conditions, for that assay. Gray indicates undetected RNA for that condition. **G.** Fraction of DORC genes (*n*=1,128) overlapping genes previously linked to super-enhancer regions under different cellular contexts (*n*=86). Only the top 50 gene sets are shown. **H.** Cumulative fraction of super-enhancer associated genes overlapping DORC genes with the addition of each cellular context

**Figure S5 (related to Figure 4).**
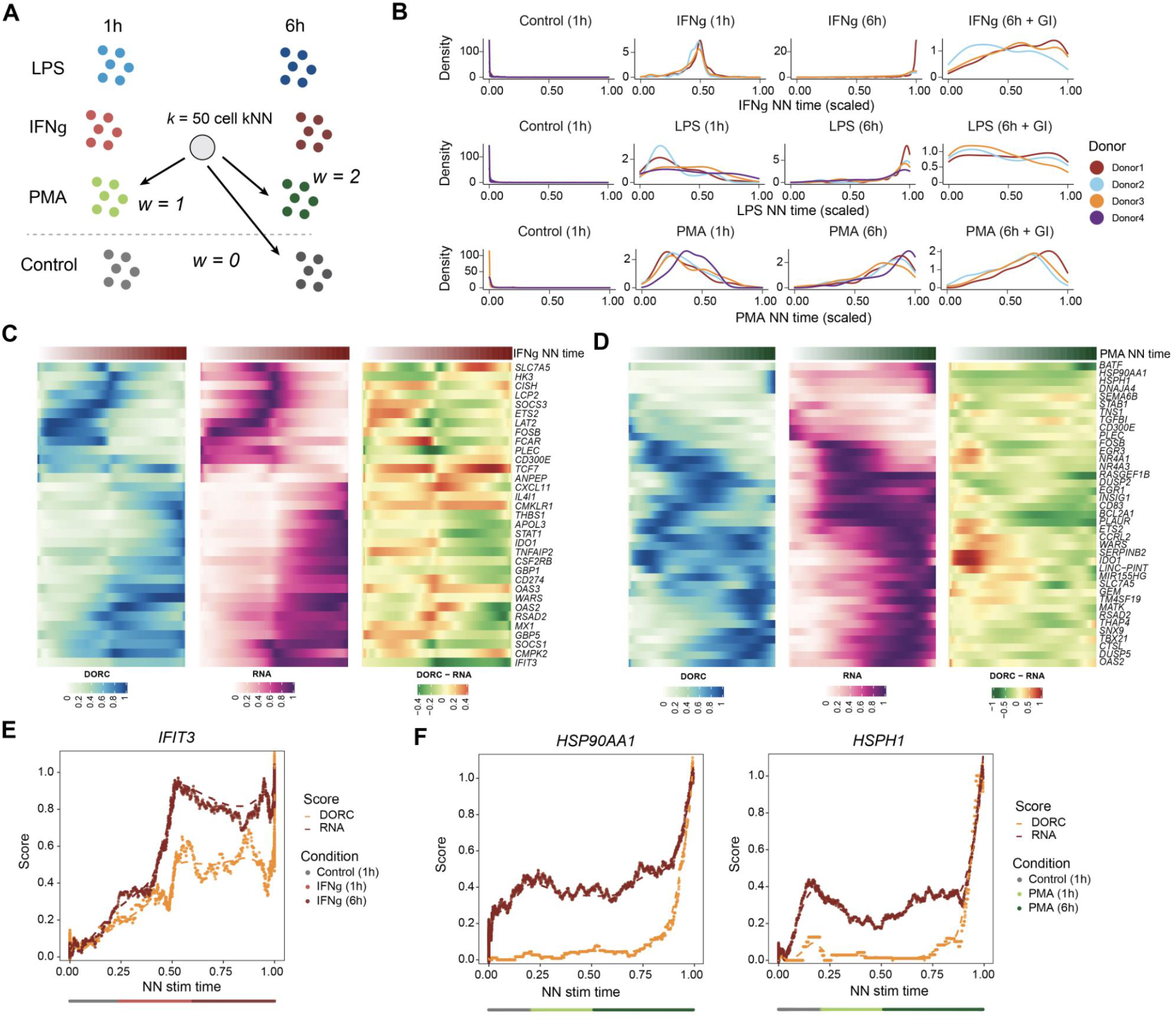
**A.** Schematic of stimulation nearest neighbor (NN) stimulation time estimation using the weighted average of cell k-nearest neighbors (kNN), per condition. **B.** Cell density distributions of NN stimulation time shown for scATAC monocytes **C.** Heatmaps highlighting smoothed normalized DORC accessibility, RNA expression and residual (DORC - RNA) levels for DORC genes with respect to IFNγ NN stimulation time for Control 1h and IFNγ 1h/6h monocyte cells (related to Fig 4C). **D.** Same as in C., but for PMA NN stimulation time-associated DORCs in monocytes. **E.** Same as in Fig 4D, but for IFNγ NN stimulation time in monocytes. **F.** Same as in E., but for heat shock protein encoding genes *HSP90AA1* and *HSPH1* with respect to PMA NN stim time.

**Figure S6 (related to Figure 5).**
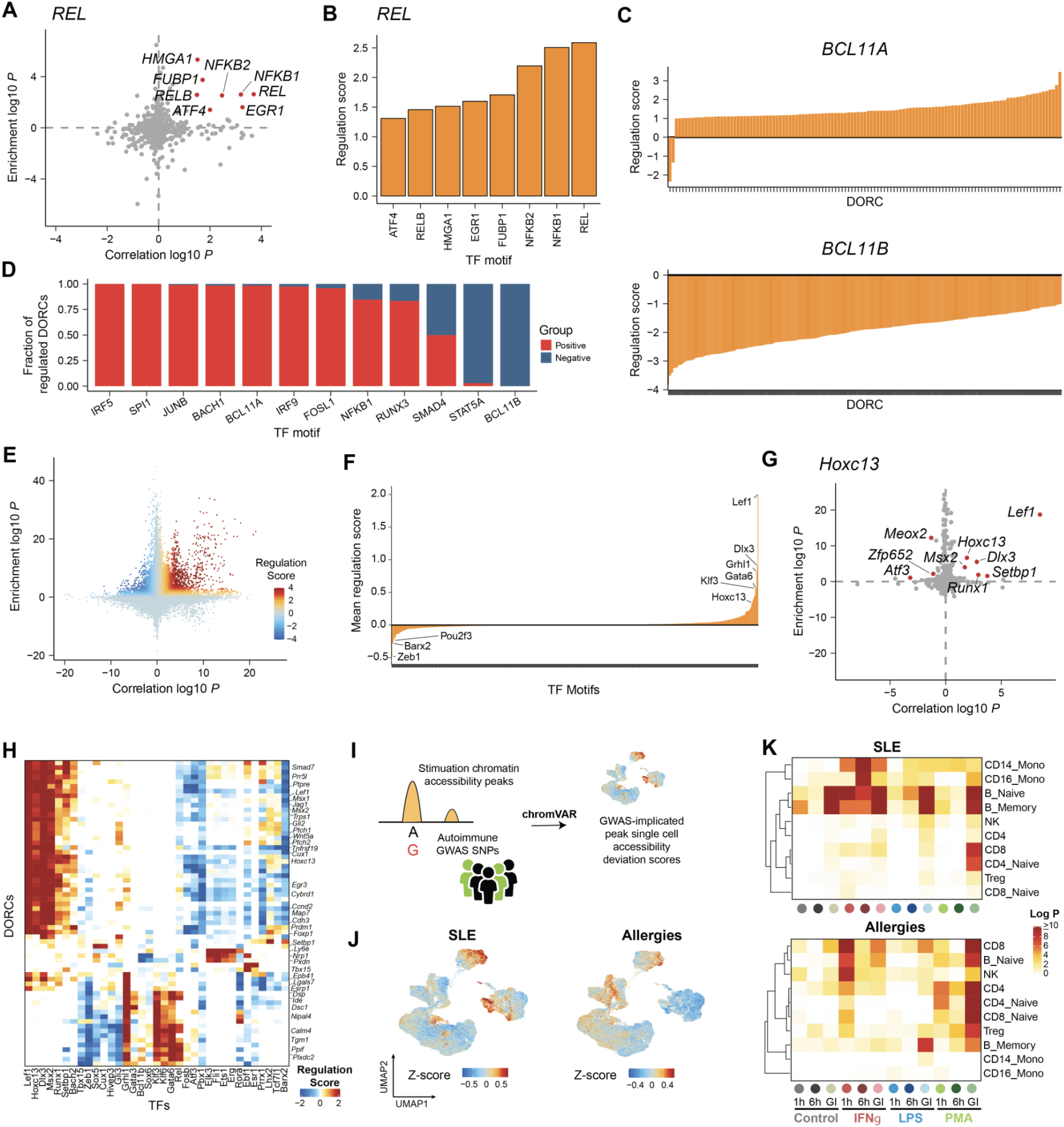
**A.** Scatter plot of TF motif enrichment among DORC KNN peaks versus TF RNA expression correlation with DORC accessibility for DORC *REL*. Red points indicate candidate drivers of *REL* (absolute regulation score ≥ 1) **B.** Regulation scores for candidate drivers of *REL* shown in A. **C.** Regulation score distributions for all putative DORCs (absolute regulation score ≥ 1) driven by *BCL11A* (top) *or BCL11B* (bottom). **D.** Barplot showing fraction of positively vs negatively associated DORCs for each TF highlighted in H. **E.** Scatter plot of all TF to DORC associations generated using FigR, run using mouse skin SHARE-seq data. **F.** Mean regulation scores highlighting main TF activators and repressors (analogous to 5E), corresponding to associations shown in E. **G.** Same as in A, but for mouse SHARE-seq derived associations for DORC *Hoxc13*. **H.** Heatmap of regulation scores between top TFs and a filtered subset of previously determined DORCs (absolute(regulation score) ≥ 2). **I.** Overview of our approach to integrate GWAS variants for 14 autoimmune/inflammatory diseases with stimulation scATAC-seq data. **J.** UMAP of scATAC-seq cells (n=62,219) colored by accessibility Z-scores based on peak-SNP overlaps for Systemic Lupus Erythematosus (SLE) and Allergies GWAS variants. Scores shown are smoothed among *k*=50 cell nearest neighbors, and thresholded at +/- 3 s.d. for visualization. **K.** Heatmap of significance estimates for peak-SNP overlap Z-score combined per condition and per cell type using Fisher’s method, shown for SLE and Allergies GWAS variants.

## Notes

https://github.com/buenrostrolab/stimATAC_analyses_code

https://buenrostrolab.shinyapps.io/stimFigR/

